# JiangShi: a widely distributed Mucin-like protein essential for Drosophila development

**DOI:** 10.1101/2022.04.20.488880

**Authors:** Yueping Huang, LingLing Li, Yikang S. Rong

## Abstract

Epithelia exposed to elements of the environment are protected by a mucus barrier in mammals. This barrier also serves to lubricate during organ movements and to mediate substance exchanges between the environmental milieu and internal organs. A major component of the mucus barrier is a class of glycosylated proteins called Mucin. Mucin and mucin-related proteins are widely present in the animal kingdom. Mucin mis-regulation has been reported in many diseases such as cancers and ones involving the digestive and respiratory tracts. Although the biophysical properties of isolated Mucins have been extensively studied, *in vivo* models remain scarce for the study of their functions and regulations. Here we characterize the Mucin-like JiangShi (JS) protein and its mutations in the fruit fly Drosophila. JS is an extracellular glycoprotein with domain features reminiscent of mammalian non-membranous Mucins, and one of the most widely distributed Mucin-like proteins studied in Drosophila. Both loss and over-production of JS lead to terminal defects in adult structures and organismal death. Although the physiological function of JS remains poorly defined, we present a genetically tractable model system for the *in vivo* studies of Mucin-like molecules.

## Introduction

Epithelial surfaces in animals are in contact with the environment. These surfaces are present in many places, including the respiratory, digestive and reproductive tracts, as well as body cavities such as the ear, eye, and mouth. In mammals, these surfaces are known to be protected by the mucus barrier or mucosal barrier. The mucus barrier serves a physical, chemical and an immunological role of protection. It also lubricates, such as the role of tears, and mediates substance exchanges (gas, water etc.) between the environment and internal organs. A major component of the mucus barrier is a class of heavily glycosylated proteins called Mucin. Mucins are commonly mis-regulated in human diseases and their abnormal presence has been used as biomarkers for disease diagnosis (reviewed in Ballester et al. 2019; Bhatia et al. 2019; Hannson 2019; Marimuthu et al. 2021). Therefore, a better understanding of how Mucin’s functions are regulated bears medical significance.

Mucins are large in number, with human having more than 20 different molecular classes (reviewed in Bansil et al. 2018; Wagner et al. 2018). In addition, Mucin molecules can be quite large, some of them having well over a few thousand amino acid residues (reviewed in Perez-Vilar and Hill 1999; Dekker et al. 2002; Thornton et al. 2008). Mucins are abundantly present and some of them have been effectively purified for in-depth studies of their biophysical properties (e.g., Carlstedt et al. 1983; Sheehan and Carlstedt 1984; Carlstedt et al. 1995). On the other hand, there is a limited number of *in vivo* models for the study of Mucin regulation, with the mouse being the overwhelming choice of model (e.g., Spicer et al. 1995; Velcich et al. 2002; Heazlewood et al. 2008; Roy et al. 2014). In 2014, a model was established for monitoring Mucin physiology in live zebra fish (Jevtov et al. 2014). Therefore, more *in vivo* models are needed, particularly ones with facile genetics.

Mucins do not share high degree of conservation at the level of primary amino acid sequence. However, different classes of Mucins share unique molecular architectures that are conserved. For example, for non-membrane associated Mucins, the N- and C-termini are rich in Cysteine residues that mediate inter-molecular cross-linking between mucin monomers. The central part of these Mucins is rich with Proline, Threonine and Serine residues, making up the PTS-rich domain. Mucins are heavily glycosylated molecules, with some Mucins having 80% of their weight coming from sugar molecules. The Thr and Ser residues in the PTS-rich domain are sites of O-linked glycosylation. Mucins often show N-linked glycosylation as well. Based on these limited but conserved features of Mucins, mucin-like molecules have been identified in a large variety of organisms, including insects (Lang et al. 2007; Schwientek et al. 2007; Syed et al. 2008; Dias et al. 2018), opening up the possibility of introducing new models for Mucin studies.

Mucin-like molecules in Drosophila have been preliminarily characterized. In addition, loss of function studies have been performed on several Mucin-like proteins in Drosophila and other insects (e.g., Garfinkel et al. 1983; Wilkin et al. 2000; Korayem et al. 2004; Syed et al. 2012; Reis et al. 2016; Lou et al. 2019; Kim et al. 2020). Nevertheless, the studied molecules so far have either a narrow or unclear tissue distributions that might not be a good representation of general Mucins. In this study, we identified the JiangShi (JS) protein, encoded by the previously uncharacterized *CG14880* gene, as a novel mucin-like molecule broadly produced during Drosophila development. JS has a typical “Cys-PTS-Cys” organization of Mucins. It is a secreted glycoprotein, and distributes at places where an epithelium-environment interphase exists, consistent with its proposed function as a component of the mucus barrier. Homozygous *js* mutant animals survive to adulthood but die right after eclosion suffering from leakage of bodily fluid from ruptured legs. Interestingly, overexpression of the wildtype protein and its derivatives exerts a dominant negative effect on the endogenous JS functions. JS homologs are readily identified in other insects and crustaceans. We thus identified an additional Mucin-like molecule essential for the development of an insect, and established a promising genetic model for the study of Mucins.

## Materials and Methods

### Drosophila stocks and crosses

Flies were raised on standard cornmeal media and kept at 25°C. The *w^1118^* stock was used as a control. The *js^EY^* stock (BL#16605), a chromosomal deficiency of the *js* locus (*Df(3R)Exel6269*, BL#7736), a *P* transposase insertion at 99B (BL#3664) and a multiply marked chromosome *3* (BL#1783) were obtained from the Bloomington Stock Center of Indiana USA.

#### Cas9-mediated mutagenesis of js

Cas9 induced *js* mutations were obtained using an approach in which the Cas9 enzyme is expressed from the *vasa* promoter in a transgene and the gRNA from the *U6* promoter in another transgene. The gRNA-targeted sequence at the ATG codon of JS is 5’-GAGAACAATCGTCGAAATGT**TGG** (sgRNA-N), and 5’-GGAATCAGGACTATCTGCGC**AGG** (sgRNA-C) at the STOP codon (the PAM sequences in bold). All mutations were verified by PCR amplification followed by sequencing using DNA from homozygous flies as templates. The mutant sequences are provided in Sequencing Results in Supplemental Materials.

#### Mobilization of the P element in js^1^

The *js^1^* allele is associated with a *white^+^*-marked *P* element, which produces pigmented eyes in an otherwise *white*-mutant background. We introduced a third chromosome transposase source (*⊗2-3*) into *js^1^* heterozygous flies and recovered white-eyed progeny carrying the original *js^1^* chromosome *3*. These chromosomes were made homozygous and checked for the status of the recessive js phenotype. Multiple independent excision events (>5) that restored the js phenotype were sequenced and all had precise excision of the P element. Several white-eyed events retain the recessive js phenotype. They were subject to genomic PCR aimed at amplifying across the insertional site of the original P element. One such allele, *js^3-1^*, yielded a PCR product of about 7.5kb. Sequencing revealed the presence of a part of the original P element (sequences are provided in Sequencing Results in Supplemental Materials.). The other events did not yield PCR products presumably because the larger size of the remaining P element fragment making the PCR inefficient.

#### Complementation tests of js alleles

To map the original *js^1^* allele, a multiply marked third chromosome was used to map the *white^+^* marker between *cu* and *e*, and close to *Sb* subsequently. Chromosome deficiencies of the regions were used in complementation tests further narrowing the range to about ten genes. Genomic PCR with primers one kb apart covering the entire region was performed using DNA from *js^1^* homozygotes as template. Elongation time was limited so that a large insertion between the primers, i.e. the *P* element, would not yield a PCR product. This PCR test tentatively pinpointed the P element insertion site at the first exon of *CG14880*, which was later confirmed by PCR and sequencing of imprecise excision events induced by P transposase.

To conduct pairwise complementation tests amongst *js* alleles (*js^1^*, *js^3-1^*, *js^EY^*, *js^cas-1^*, *js^cas-2^*, *js^c-term^*, *df*), 5-8 pairs of *js* heterozygotes carrying two different alleles were mated and transferred to a new vial after five days. They were discarded 10 days after they were initially crossed. Progeny were scored for 10 days after they started to emerge. Typically, more than 200 progenies emerged from each cross and no survivors were *js* mutants. Between 10-60% of the *js* mutants did eclose but soon died and displayed the js phenotype (for details see Results), and they were not counted as survivors.

### Plasmid construction

For Cas9 mediated tagging at the C-terminus of JS, a ∼2kb fragment centered at the STOP codon (from 948 bp 5’ of “T” of the TAG codon to 1103 bp 3’ of “G”) was subcloned from genomic DNA and a DNA fragment encoding the three fluorescent proteins (mCherry, dsRED and EGFP) was individually inserted just upstream of the STOP codon using bacterial recombineering (Zhang et al. 2014). These plasmids served as donor DNA in homologous recombination in which the two DNA fragments flanking the fluorescent gene are homologous to each side of the Cas9-induced DNA break on the chromosome. This donor plasmid was injected into flies expressing Cas9 and a gRNA targeting the STOP codon of *js* (sgRNA-C). Knock-in events were screened by PCR and verified by genomic PCR followed by sequencing. Sequences are provided in Sequencing Results in Supplemental Materials.

For JS overexpression, full-length *js* cDNA was amplified from cDNAs reverse transcribed from RNA isolated from wildtype adults. The cDNA was cloned into the multiple cloning sites of pUASTattB generating the *UAS-js* construct for subsequent phiC31-mediated integration at the 75B genomic location. A fragment encoding dsRED was inserted into *UAS-js* just before the STOP codon of JS by recombineering generating the *UAS-js-dsRED* construct. Based on *UAS-js-dsRED*, two small deletions were introduced individually by site-directed mutagenesis generating the *UAS-⊗CBD-I* and -*II* constructs. All constructs were confirmed by sequencing. Sequences are provided in Sequencing Results in Supplemental Materials.

### Western blot reagents and assays

A corresponding DNA fragment encoding the JS antigen was subcloned from cDNA into pET28a for expression. Bacterial expression was induced with 0.1 mM IPTG, and the recombinant protein was purified as inclusion bodies. Briefly, inclusion bodies were separated from soluble fractions by centrifugation. Pellets were washed three times with 2M Urea and 1% Triton X-100, and three times with 2M Urea alone. Each wash constituted a brief sonication of the pellet in wash buffer followed by centrifugation and disposal of the supernatant. The washed pellet was dissolved in 8M Urea and used in immunization of rabbits to generate polyclonal anti-sera. Rabbit anti-sera were used at a dilution of 1:5000 for Western blotting. The transfer of JS protein onto PVDF membrane was done by a protocol developed for studying Mucin-D (Kramerov et al. 1996 and 1997). Briefly, extracts were prepared with traditional SDS loading buffer. SDS-PAGE was run with the standard 1XTGS (Tris-Glycine-SDS) buffer at a Voltage of 100V, and transferred in 10mM sodium borate buffer (pH9.2) for 3h at 0.4A on ice. The WGA assay was performed as described (Fujioka et al. 2018), except using the transferring conditions described above.

### Microscopy

Life imaging of fluorescently labelled JS proteins produced from *js* knock-in alleles was performed on a Zeiss Image2 fluorescence microscope. Embryos were mounted in 50% glycerol and observed directly. Larvae were mounted in 50% glycerol, heated on a 65°C heating block for a few seconds till they stopped moving, and covered with a coverslip for microscope observation. We confirmed that while this heat-induced immobilization facilitates life imaging, it does not alter the expression pattern nor intensity of the fluorescently tagged JS proteins. Pupae and adults were mounted in glycerol and observed directly.

For immunostaining, anti-mCherry antibody from Abcam (ab167453) and anti-dsRED from Santa Cruz (sc-101526) were used at various dilutions but did not produce satisfactory results.

Transmission and scanning EM analyses were performed at the core facility of the School of Life Sciences Sun Yat-sen University China, following standard protocols. For TEM analyses, wildtype (*w^1118^*) and *js* mutant tissues were prepared as follows: legs were dissected from adults within one day after eclosion. Gut, pericardial cells and trachea were dissected from larvae in 3rd instar. Tissues were fixed overnight in 4% glutaraldehyde at 4°C. After three washes with 0.1 M sodium cacodylate (pH 7.2), tissues were stained with 1% osmium tetroxide for 1hr at room temperature. They were washed three times again, and stained with uranyl acetate overnight. After a standard ethanol dehydration series (5 minutes each in 25%, 50%, 75%, 95% EtOH, and 3x10 minutes in 100% anhydrous ETOH), tissues were rinsed in propylene oxide twice before they were embedded using standard procedures. Thin sections (100 nm) were cut and collected on support grids, and stained with uranyl acetate for 15 min, followed with 10 min in lead citrate. Micrographs were taken at 120 kV on a JEM-1400 TEM microscope. For SEM analyses, legs were dissected from adults within one day after eclosion, fixed for overnight in 4% glutaraldehyde at 4°C, washed three washes with PBS (pH 7.0), dehydrated with a graded ethanol series (5 minutes each in 25%, 50%, 75%, 95% EtOH, and 3x10 minutes in 100% anhydrous ETOH). They underwent the processes of critical point drying and metallizing before micrographs were taken on a Hitachi S-3400N microscope.

### Silver nitrate feeding

Feeding was performed as described previously (Zhang et al. 2013a). Briefly, first instar larvae of *js^1/1^* and *js^1^/TM6B* were selected, placed separately on agar-only plates supplemented with regular yeast paste or yeast paste containing AgNO3 (2.0 g yeast in 3.5 ml 0.005% AgNO3 solution) and allowed to develop at 25°C until adulthood. Pupae from both genotypes were skinned for photography.

## Results

### The Jingshi phenotype

We fortuitously recovered a recessive mutation on the third chromosome that can only be kept as a heterozygous stock, and homozygotes display the following novel phenotypes. They develop into pharate adults and the majority eclose successfully. However, adults emerged with weak legs and showed uncoordinated movements, hence the Chinese name *jiangshi* (*js*): a walking dead. Leg joints on all of the adults carry dark colored and sticky substances (Figure 1A). These adults ultimately drowned in food or were stuck to the side of the vials, likely due to the blackish substances coming from the joints. The description of the *jiangshi* phenotype matches very well with that of a previously isolated but nonextant *arthritics* (*arth*) mutation. Without a better description of the *js* phenotype, we use that for the *arth* mutation verbatim: “legs weak with pigmented joints; tarsal segments frequently askew with claws fused; movements somewhat uncoordinated; brownish-black pigment present at joints …; most frequently in meso- and metathoracic legs between femur and tibia but sometimes between coxa and trochanter or proximal to coxa.” (flybase.org). Since *arth* has been mapped to the *X* chromosome (Fleming et al. 1989), *arth* and *js* are two different genes.

**Figure 1.**
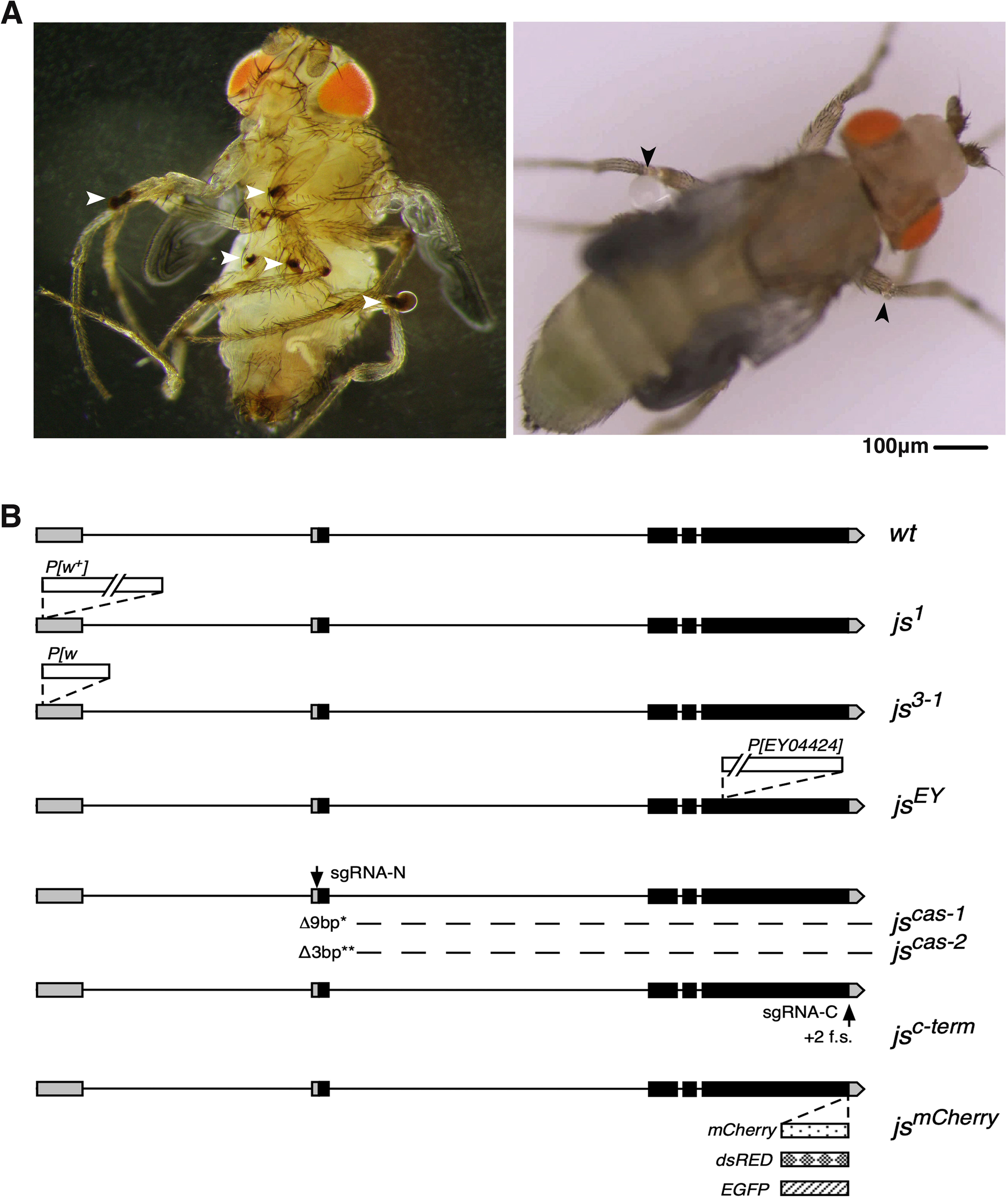
The phenotype and genomic structure of *js* alleles. **A.** Morphological defects of *js* mutant adults. At the left is a darkfield image of a *js^1/df^* adult showing blackish substance at multiple leg-joints indicated with white arrowheads. At the right is a brightfield image showing liquid droplets forming at the leg-joints (arrowheads) of a *js^1/df^* adult. **B**. Diagrams of the *js* locus. The genomic structure of the wildtype (*wt*) *js* locus is shown at the top with exons shown as boxes and introns as lines. Non-coding exons are shaded in grey and coding ones in black. The arrow indicates the direction of transcription. The *js^1^* allele has a *white^+^* (*w^+^*) marked *P* element inserted at the first but non-coding exon so that *js^1^* flies have pigmented eyes. The *js^3-1^* allele has a ∼7kb fragment of the original *P* element remained in which a part of the *w^+^* gene was deleted so that *js^3-1^* flies are white-eyed. The *js^EY^* allele has a *P* element inserted in the largest coding exon of *js*. The *js^cas-1^* (a 9 bp deletion) and *js^cas-2^* (a 3 bp deletion) mutations were generated by CRISPR/Cas9 mediated mutagenesis utilizing the “sgRNA-N” guide RNA, targeting the START codon of *js* (marked with an arrow). The *js^c-term^* alleles are +2 frame shift (f.s.) mutations induced by CRISPR/Cas9 utilizing the “sgRNA-C” guide RNA, marked with an arrow. The *js^mCherry^* knock-in allele has a DNA fragment encoding mCherry inserted at the endogenous *js* locus just upstream of the STOP codon. Similarly, a fragment encoding dsRED or EGFP was knocked into *js* at the identical position.

A video recording the eclosion of a *js* adult (Supplemental Materials Movie S1) clearly shows that fluids form droplets of various sizes on leg joints after the fly emerges from the pupal case (see also Figure 1A). We therefore postulate that leg joints of *js* adults rupture during the strenuous eclosion process, letting out body fluids that form the basis for the blackish substances. The change from colorless droplets to blackish substances might be related to the melanization process of Drosophila hemolymph. We did not further investigate the nature of the black matter but chose instead to focus on uncovering the genetic cause for this unique phenotype.

### The *js* gene

The original *js* allele, which we name *js^1^*, is associated with a *white^+^* eye marker. Using a combination of recombination-based and chromosomal deficiency-based mapping methods, followed by PCR and genomic sequencing, we determined that *js^1^* is associated with a *P* transposable element inserted into the 5’ *UTR* of the gene *CG14880* (for mapping details see Materials and Methods). Figure 1B shows the genomic structure of the *js* locus and its various alleles. Several lines of evidence support that mutations in *CG14880* cause the *js* phenotype. First, we mobilized the *P* element in the germline by expressing *P* transposase, and recovered events of precise excision of the element that reverses the mutant phenotype (see Materials and Methods). We also recovered imprecise excision events losing only part of the *P* element but retaining the phenotypes (the *js^3-1^* allele). Secondly, an independent *P* element insertion into one of the coding exons of *CG14880*, causes the recessive js phenotype (the *js^EY^* allele). Thirdly, we recovered two CRISPR/Cas9-induced small deletions of the start code of *CG14880*. These mutations also cause the js phenotype (the *js^cas-1^* and *js^cas-2^* alleles). Fourthly, we used CRISPR/Cas9 to make mutations around the Stop codon of *CG14880*, and recovered multiple alleles deleting the Stop codon while causing a +2 frameshift, which displayed the js phenotype (the *js^C-term^* allele). Lastly, all of the above point mutations of *js*, when trans-heterozygous with each other or with a chromosomal deficiency that deletes the *CG14880* region (*Df(3R)Exel6269*), produced the js phenotype (see Materials and Methods for details of the complementation tests). Therefore, our extensive genetic analyses establish that mutations in *CG14880* are responsible for the js phenotype.

### JS protein has features of Mucins

The *js* gene encodes a protein of 637 residues (Figure 2A). The first 30 residues of JS are predicted to be a signal peptide suggesting that JS is a secreted protein (Supplemental Materials Figure S1). C-terminal to the signal peptide, starting around the 51^st^ residue, is a domain of about 60 residues that has been annotated as a Chitin Binding Domain (CBD). The CBD domain is highly enriched with conserved Cysteine residues (∼10%). Therefore, mature JS protein has a Cys-rich N-terminus. At the very C-terminus of JS, we identified a conserved motif (about 20 residues in length) also enriched with Cysteines (∼20%), which we named Cysteine Rich Motif (CRM). Between CBD and CRM lies the majority of the JS residues, which are noticeably enriched with Proline, Threonine and Serine residues (PTS-rich): 175 of the 510 intervening residues (34%). PTS domains of mammalian Mucins are modified with O-linked glycosylation, while the presence of N-linked glycosylation is also common. In JS, two N-linked glycosylation sites have been identified (Baycin-Hizal et al. 2011), with one inside CBD (Figure 2A). The above features that we described for JS: a secreted protein with two Cys-rich domains flanking a PTS-rich region, are consistent with the basic domain organization of non-membranous Mucin-like molecules previously identified in various vertebrates. However, we note that JS was not identified in prior bioinformatic studies of insect Mucin-like molecules (e.g., Lang et al. 2007; Syed et al. 2008). We note that clear JS homologs based on sequence homology are present in insects and crustaceans (Figure 2A, Supplemental Materials Figure S2).

**Figure 2.**
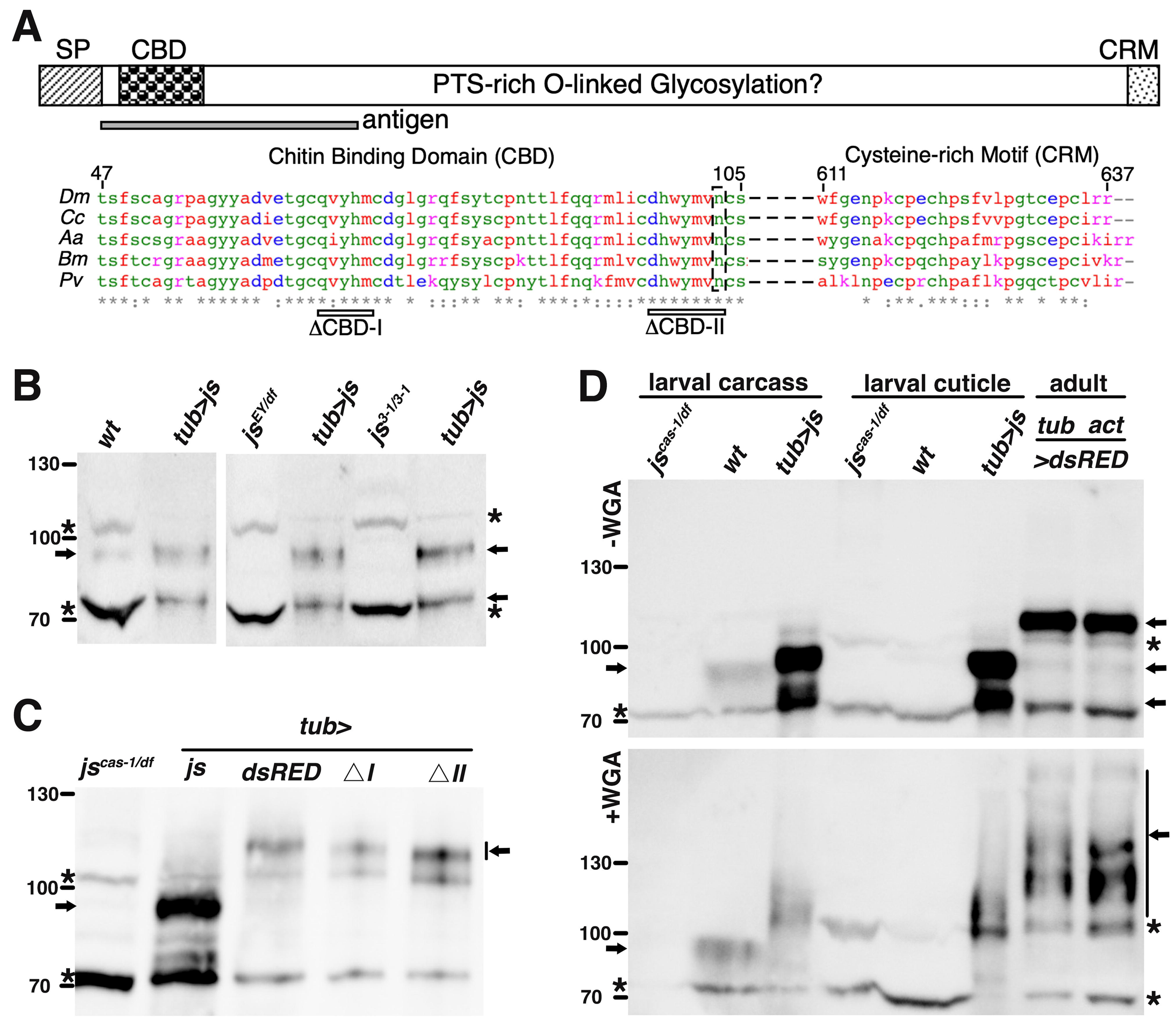
Features of the JS protein and the detection of its glycosylation. **A**. A diagram of the structural elements of the JS protein is presented at the top: SP (signal peptide); CBD (chitin binding domain); Proline Threonine Serine (PTS)-rich domain; and CRM (Cysteine rich motif). The antigen used for raising antibodies is indicated beneath the domain diagram. Sequence alignments, generated by clustal omega, for the CBD and CRM domains are shown for the following species: *Dm* (*Drosophila melanogaster*, NP_650538.1), *Cc* (*Ceratis capitata*, XP_012157262.1), *Aa* (*Aedes aegypti*, EAT42245.1), *Bm* (*Bombyx mori*, XP_004927149.1), and *Pv* (*Penaeus vannamei*, pacific white shrimp, XP_027211762.1). At the top of the alignment, the residue numbers are provided for the Drosophila protein. Beneath the alignment, the ranges of the two CBD deletions are shown for I (deletion of 5 residues) and II (deletion of 7 residues). The N-linked glycosylation site is marked with a black box. **B**. Western blot detection of JS protein in adult tissues. Whole extracts from wildtype (*wt*), two *js* mutants and adults overexpressing full-length JS from the *tubulin* Gal4 driver (*tub>js*) were used for Western blotting. The overexpressing extracts were loaded at a 1:50 dilution. The JS protein bands are indicated with arrows. Non-specific bands, marked with asterisks, were used as loading controls. Note that one of the JS bands (∼70KD) in the overexpression extracts ran close to one of the major non-specific bands. Markers, with sizes indicated in KD, are shown to the left. **C**. Western blot detection in adults overexpressing various forms of the JS protein. Extracts from flies overexpressing various derivatives of the JS protein were used along with extracts from *js* mutant extract. The overexpression extracts were used at a 1:50 dilution. Overexpression was driven by the tubulin Gal4 driver (tub>). *js*: full-length JS overexpression; *dsRED*: full-length JS tagged with dsRED at the C-terminus; *⊗I*: CBD-I deleted JS tagged with dsRED at the C-terminus; *⊗II*: CBD-II deleted JS tagged with dsRED at the C-terminus. The JS protein bands are indicated with arrows. Non-specific bands are marked with asterisks. Markers, with sizes indicated in KD, are shown to the left. **D**. Western blot-based WGA assay for detection of JS protein and its glycosylation state. Extracts were derived from larval cuticles (middle three lanes), larval carcasses devoid of cuticles (left three lanes) and whole adults (right two lanes). Larval extracts were from *js* mutants, wildtype, or animals overexpressing full-length JS driven by tubulin Gal4 (*tub>js*). Adult extracts were from animals overexpressing full-length JS tagged with dsRED, driven by tubulin (tub) or actin5C (act) Gal4 driver. The overexpression extracts were used at a 1:10 dilution. Bands corresponding to JS are indicated by arrows, while none-specific bands with asterisks. Extracts were run in parallel on two gels, the lower of which contained WGA (+WGA). Markers, with sizes in KD, are indicated to the left.

An essential feature of Mucins is O-linked Glycosylation. To investigate whether JS is glycosylated, we generated antibodies against an antigen consisting of the first 198 residues of the mature JS protein (Figure 2A). In normal Western blots (several shown in Figure 2), we observed a band of about 95KD in size. The signal for this band is weak and variable, and it is different from JS’s predicted size of 70KD, but it is consistently missing in *js*-mutant extracts. We reasoned that the possible causes for the weak JS signals on a Western blot could be two folds that are not mutually exclusive. First, the level of JS might be relatively low. Secondly, our antibodies, raised against a bacterially expressed JS antigen, might be inefficient in recognizing the endogenous JS proteins especially considering that JS is likely a glycosylated protein with a potentially high propensity to form intra- and inter-molecular disulfide bonds. We note that the JS antigen encompasses a site where the endogenous JS protein is modified by N-linked glycosylated (Figure 2A). To gain more confidence on the specificity of our antibodies, we overexpressed JS using a construct that places *js* cDNA encoding the full-length protein as well as its various derivatives under the control of the Gal4 activator. We used either a *tubulin* (*tub*) or an *actin5C* (*act*) Gal4 driver to achieve ubiquitous overexpression. As shown in Figure 2B,2C,2D, our antibodies very strongly recognize a dominant band of about 95KD and a minor one of about 70KD specifically under the condition of JS overexpression, suggesting that both protein species are produced from the JS-overexpressing transgene and further validating our antibodies. Note that the overexpressing extracts were loaded at much-diluted amounts (1:10 to 1:50). We postulate that the size increase of the endogenous JS protein is related to JS being a putative glycoprotein.

The traditional method of using de-glycosylases (Ziv et al. 2012) proved unproductive in characterizing JS’s state of glycosylation. We employed a gel electrophoretic method that cleverly takes advantage of the ability of Wheat Germ Agglutinin (WGA) to bind and retard the mobility of glycoproteins in traditional SDS PAGE gels (Kubota et al. 2017). As shown in Figure 2D, the endogenous and particularly the over-expressed JS display mobility retardation in a WGA gel when compared with one without WGA, and when markers and non-specific bands were used as internal controls for protein mobility. Interestingly, both the 70KD and 95KD bands from the JS-overexpressed samples showed WGA-retarded mobilities, suggesting both are glycosylated but perhaps to different extents. Therefore, how the two protein species differ in mobility cannot simply be due to the absence or presence of glycosylation. Interestingly, Overexpressed JS is abundantly present in extracts made from cuticles of third instar larvae (Figure 2D), consistent with JS being a secreted protein. In summary, JS protein bears common features of Mucin-like molecules with the important presence of glycosylation.

We note that WGA is often used for the detection of N-acetyl-D-glucosamine (O-GlcNAc), while mammalian mucins are enriched with N-acetyl-galactosamine (O-GalNAc). Therefore, although our results support JS being a glycoprotein, whether it possesses O-GalNAc modifications requires further investigations.

### JS is a secreted protein highly enriched at epithelia and widely expressed in development

Although our antibodies were able to recognize the JS protein on Western blots, they failed to generate satisfactory results in immunostaining experiments using JS non-overexpressing tissues. To study the localization of endogenous JS, we generated a knock-in allele of *js* by CRIPSR/Cas9-mediated homologous recombination, in which a DNA fragment encoding the mCherry fluorescent protein was placed in-frame before the STOP codon (*js^mCherry^*, Figure 1B). The mCherry tag does not disrupt JS functions as *js^mCherry^* homozygous flies are viable, fertile, and kept as a homozygous stock. Thus, the fluorescence from mCherry allows us to deduce the normal localization of endogenous JS proteins. To minimize interference from auto-fluorescence, we only identify red signals as originated from the JS-mCherry protein when the same region does not emit green/yellow fluorescence, and when the same region does not emit red fluorescence under the wildtype background (Supplemental Materials Figure S3). We present below, to the best of our abilities, a comprehensive description of JS-mCherry distribution in development.

#### Early developmental stages

The mCherry signal is first visible around twelve hours after egg laying but become prominent in first instar larvae (Figure 3A), showing prominent fluorescence at the mouthpart, pharynx, and the pericardial cells. Interestingly, right after hatching, JS-mCherry signals are visible throughout the larval cuticle and in the form of puncta (Figure 3F).

**Figure 3.**
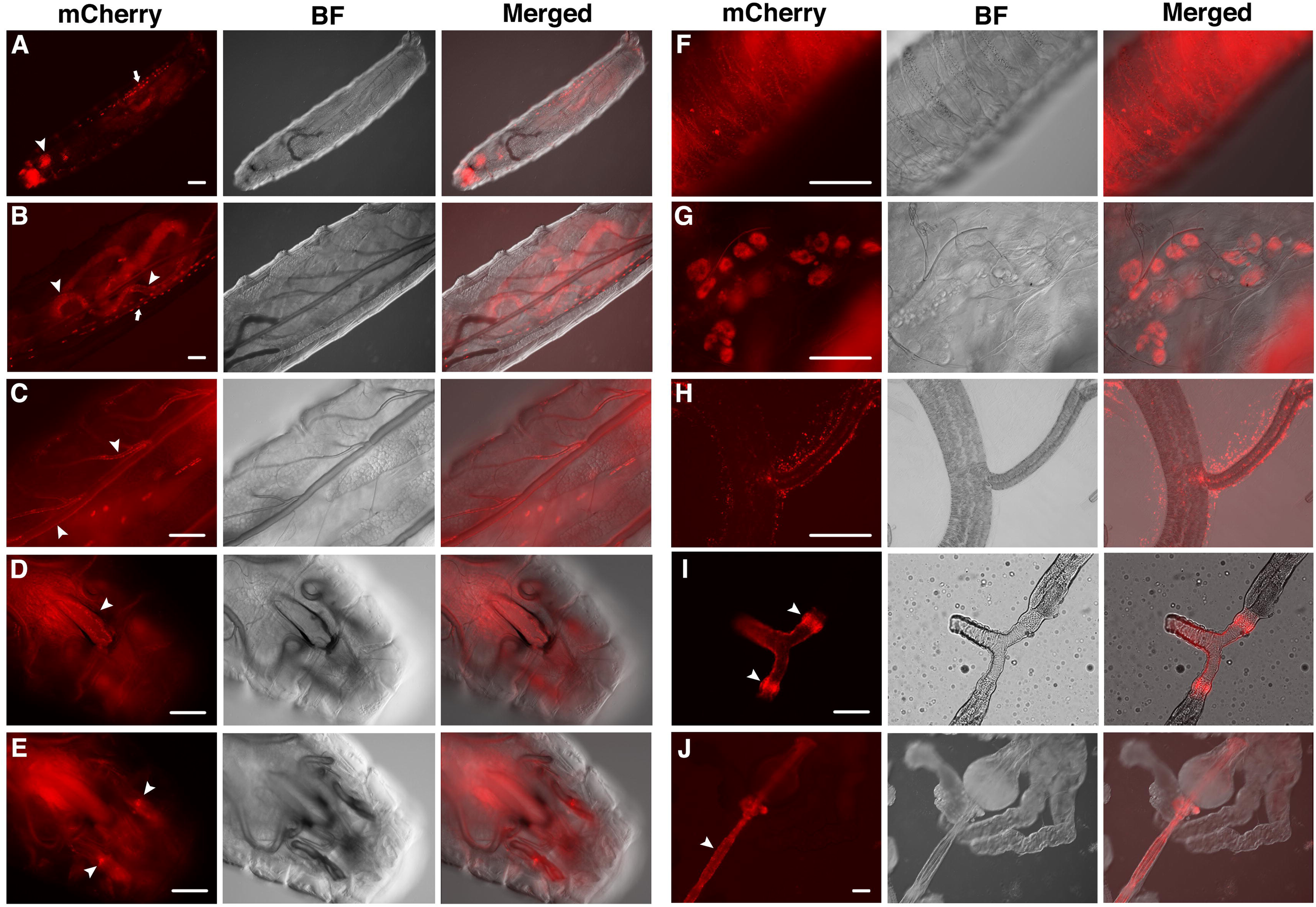
JS localization during larval development. Live images are provided as triplets: red fluorescence (mCherry), brightfield (BF) and the merged of the two. **A**, a whole mount first instar larva. Arrow indicates pericardial cells, and arrowhead indicates the mouth area. **B**, a second instar larva. Arrow indicates pericardial cells, and arrowheads indicate guts. **C**, a third instar larva. Arrowheads indicate trachea. **D** and **E**, mouth area of a third instar larva. Arrowhead in **D** indicates the pharynx. Those in **E** indicate the humerus discs. **F**, cuticular view of a first instar larva. **G**, garland cells from a third instar larva. **H**, JS puncta surrounding trachea from a third instar larva. **I**, duct area of a dissected salivary gland from a third instar larva. Arrowheads indicate the imaginal rings. **J**, dissected foregut (arrowhead) from a third instar larva. Scale bars indicate 100μm.

JS distribution does not change significantly as the larva grows. JS-mCherry signal is present at: guts (Figure 3B, J), trachea (Figure 3C, H), pharynx (Figure 3D), humerus discs (Figure 3E), pericardial and garland cells (Figure 3B, G). We observe JS signals surrounding the individual ducts and imaginal rings of the salivary glands (Figure 3I).

#### JS-mCherry molecules form puncta

We notice that JS-mCherry signals appear as puncta in various tissues (Figures 3 and 4). This focused appearance seems to be inconsistent with the expected distribution of Mucin-like proteins, which would be a more even distribution over the entire epithelium. However, one cannot rule out that these puncta only represent some of the JS molecules since our detection method depends on mCherry’s ability to fluoresce in the different chemical environments that JS-mCherry exists in. It is possible that some of the JS-mCherry molecules do not fluoresce. Results described in the next section lend support to this proposition.

**Figure 4.**
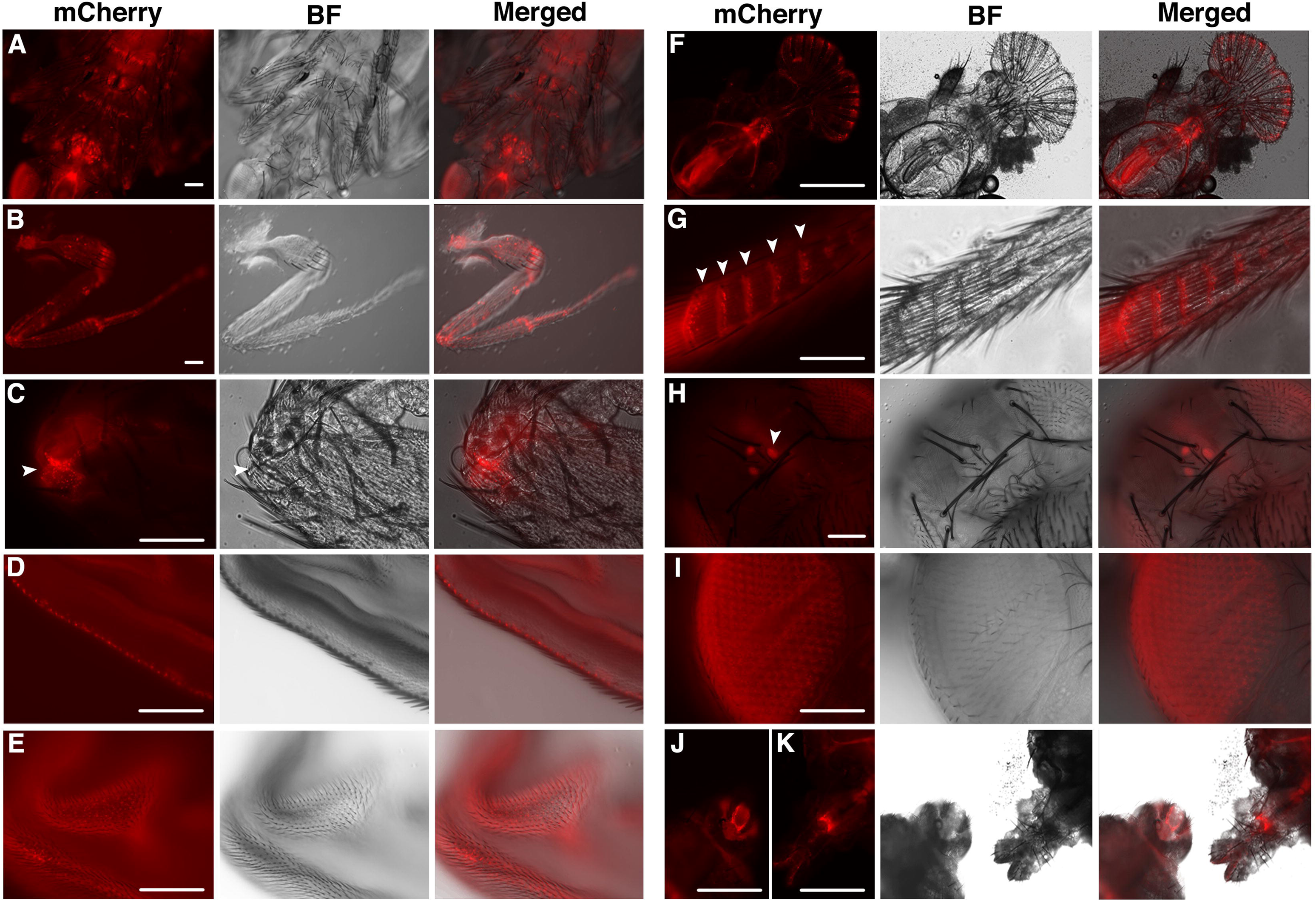
JS localization in pupal and adult stages. **A**, a late-stage pupa showing mCherry signals in multiple leg joints. **B**, a dissected front leg from an adult. **C**, a closeup picture of an adult leg joint showing JS puncta surrounding the junction (arrowhead). **D**, wing blade from an adult showing a row of JS puncta. **E**, surface view of a pupal wing showing JS puncta. **F**, mouth area of an adult. **G**, a closeup view showing rows of JS puncta (arrowheads) aligning with the rows of bristles on an adult leg. **H**, JS puncta in the three ocelli (arrowhead). **I**, surface view of an adult eye showing JS puncta surrounding each ommatidium. **J**, a female genital. **K**, a male genital. Scale bars indicate 100μm.

We generated other *js* knock-in alleles in which either a dsRED or an EGFP tag was placed at the C-terminus of JS (Figure 1B), similarly to the mCherry tag. Interestingly, none of these two alleles, although allowing homozygous flies to survive and reproduce, displays fluorescence similar in pattern or intensity to that of JS-mCherry, suggesting that the natural environment where JS resides is less permissive for dsRED or EGFP fluorescence. The same environment seems to be more permissive to mCherry fluorescence, although not necessary for all of the JS-mCherry populations. Therefore, it cannot be ruled out that JS-mCherry puncta that we observed represent one class of the JS molecules, perhaps as intermediates in the process of JS secretion or maturation, and that the functioning JS-mCherry molecules at the epithelia might not be fluorescently visible. In other words, we cannot be confident that most of the JS-mCherry molecules have been visually localized in our live analyses. Unfortunately, immunostaining with anti-mCherry and anti-dsRED antibodies using *js^mCherry^* and *js^dsRED^* tissues respectively did not yield satisfactory results.

#### Pupal and adult stages

JS-mCherry is abundant in all of the major joints of legs, consistent with the most prominent phenotype of *js* mutant adults (Figure 4A-C, G). It is also highly significant on wing (Figure 4D, E), the mouth (Figure 4F), genital of both sexes (Figure 4J, K). JS-mCherry punctum seems to be located at the base of each major bristle. This is most prominent for bristles on the legs and wings, possibly due to that those areas are under less interference from internal fluorescence (Figure 4D, E, G). Remarkably, we observed puncta of JS-mCherry surrounding each ommatidium in the eye and this pattern is reproducible from GFP fluorescence present in *js^egfp^* knock-in flies (Figure 4I and Supplemental Materials Figure S4). In ocelli of both *js^mCherry^* and *js^egfp^* flies, we also observed strong JS puncta (Figure 4H and Figure S4).

### Potential functions of JS

To shed lights on possible functions of JS, we conducted Electron Microscopy (EM) imaging of tissues where JS is present. Scanning EM reveals “extra” substances covering mutant leg joints, likely corresponding to the blackish substance under visible lights (Figure 5B). However, transmission EM imaging of cross sections of legs failed to uncover apparent structural differences between wildtype and *js* mutants (Figure 5A). Similarly, cross sectioning of guts (Figure 5C) or trachea (Figure 5D) did not reveal abnormalities in the mutant tissues.

**Figure 5.**
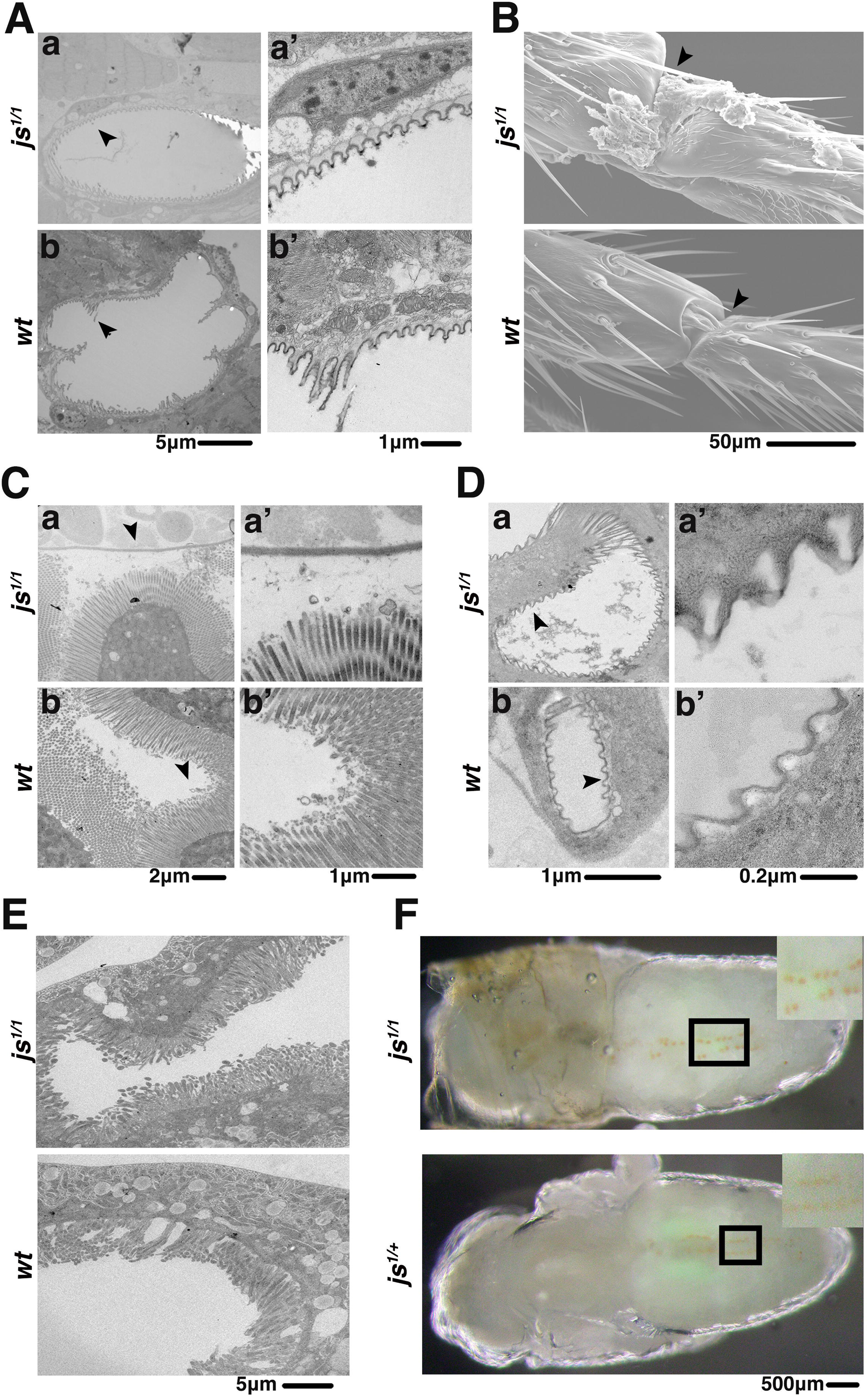
Ultrastructural studies of *js* mutants. For all images, genotypes are listed to the left. **A**. Transmission Electronic microscopic (TEM) images of leg cross-sections. The area marked with an arrow in **Aa** and **Ab** was photographed again but with a higher magnification, and shown in **Aa’** and **Ab’** respectively. **B**. Scanning EM images of leg joints, with arrowheads demarcating the area where extra substances are visible covering the joint from the mutant but not wildtype. **C**. TEM images of cross sections of larval guts. The area marked with an arrow in **Ca** and **Cb** was photographed again but with a higher magnification, and shown in **Ca’** and **Cb’** respectively. **D**. TEM images of cross sections of larval trachea. The area marked with an arrow in **Da** and **Db** was photographed again but with a higher magnification, and shown in **Da’** and **Db’** respectively. **E**. TEM images of cross sections of nephrocytes (pericardial cells). **F**. Bright field images of pupae showing the accumulation of AgNO3 in the nephrocytes, with the area marked with a box shown at a higher magnification at the top right corner.

One of the most consistent tissues where JS is present is the pericardial cells (Figure 3). Nevertheless, cross sectioning of larval pericardial cells did not reveal structural abnormalities suggestive of a functional loss in these cells (Figure 5E). Pericardial cells serve to filter the hemolymph, similar in function as mammalian nephrocytes (reviewed in Helmstadter et al. 2017). Disruption of nephrological function in Drosophila has been reported to cause sensitivity to silver poisoning (Weavers et al. 2009; Zhang et al. 2013a; Ivy et al. 2015). We therefore treated flies with Silver Nitrate (AgNO3) but did not observe an overt effect on the development of *js* mutants in that homozygous larvae fed with AgNO3 were able to eclose as adults. In addition, bright field imaging clearly shows that silver salts were successfully retained in larval nephrocytes in *js* mutant larvae (Figure 5F).

In summary, our ultra-structural studies reveal largely normal tissue morphology in *js* mutant animals leaving the potential physiological function of JS undefined.

### Overproduction disrupts JS function

Although all *js* mutant alleles behaves as recessive mutations, we discovered that overexpression of the wildtype JS protein is sufficient to produce very similar if not identical phenotypes to those in recessive mutants described previously.

In an attempt to “rescue” the js phenotypes, we constructed transgenic lines carrying a *js* cDNA clone under the control of *UAS* elements. We used various Gal4 drivers to deliver JS proteins to the *js*-mutant background. None of the tested Gal4 drivers was able to rescue the lethality of the homozygotes. In addition, we discovered that the strong and ubiquitous Tubulin-Gal4 (tub-Gal4) or Actin5C-Gal4 (act-Gal4) driven overexpression of JS in a wildtype background was sufficient to cause the js phenotypes (Figure 6A). This effect could also be reproduced by over-expressing a JS-dsRED fusion protein (Figure 6A). In addition, we constructed two separate deletions within the conserved Chitin Binding Domain (CBD), with deletion I more N-terminal to deletion II (Figures 2A), and tagged both proteins with a C-terminal dsRED moiety. Both truncated proteins were similarly overexpressed by act-Gal4 or tub-Gal4, and their effects on adult survival were measured.

**Figure 6.**
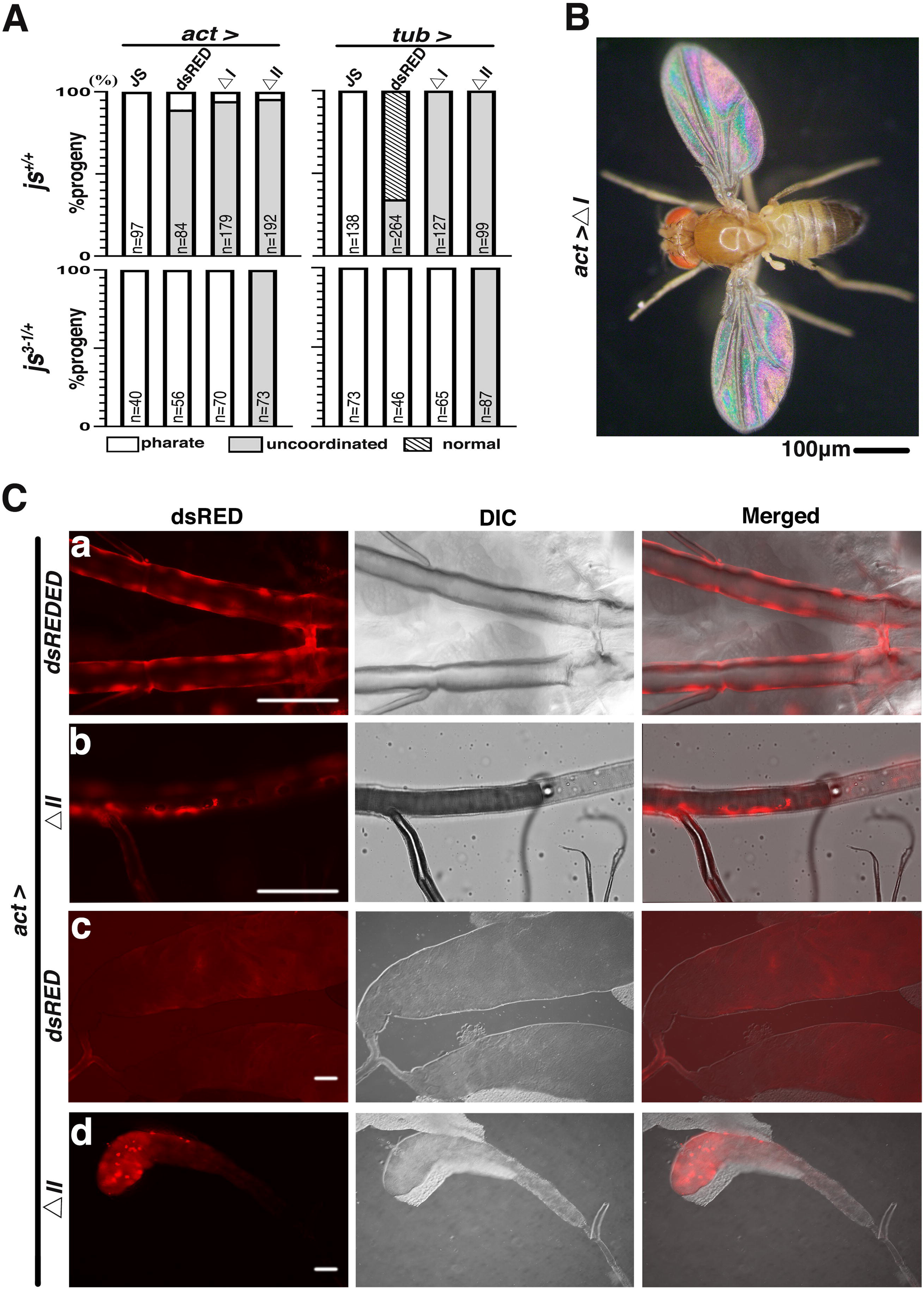
Overexpression of JS disrupts its endogenous function. **A**. Quantitative analyses of adult phenotypes from JS overexpression. Adults are classified into three classes (“pharate”, “uncoordinated” and “normal”), and their counts are plotted according to different JS over-expressing conditions. The Gal4 driver used is listed at the top with the JS proteins being over-expressed listed underneath. JS: full length JS protein; dsRED: full length JS with a C-terminal dsRED tag; ⊗I: JS protein with the CBD deletion I and a dsRED tag; ⊗II: JS protein with the CBD deletion II and a dsRED tag. The state of the endogenous *js* loci is listed to the left. The number of progenies counted are presented as “n” in each column. **B**. The “spread-wing” phenotype of an adult overexpressing ⊗CBD-I with the actin5C driver (bottom). A wildtype adult with a normal wing posture is shown at the top. **C**. Ectopic localization of dsRED-tagged JS derivatives expressed from the act-Gal4 driver in an otherwise wildtype background. Representative images are shown in triplets: red fluorescence (dsRED), DIC, and the merged image of the two. **Ca** and **Cb**, larval trachea showing big patches of dsRED fluorescence. **Cc**, dsRED localization in the main lobes of the larval salivary gland, but missing from the imaginal rings. **Cd**, aggregation of dsRED signals in secretion cells of the larval salivary gland. Scale bars indicate 100μm.

We classified JS-overexpressing adults into three phenotypic classes: pharate or short-lived adults, adults with uncoordinated movements, and normal adults (Figure 6A). Most flies, dead or alive, had black-pigmented joints on their legs. Under the wildtype (*js^+/+^*) background, act-Gal4 driven production of full-length JS proteins resulted in essentially 100% lethality (top row in Figure 6A) with most adults dying soon after eclosion and the rest died as pharate adults. Interestingly, when a C-terminally dsRED tagged JS protein was overproduced similarly, a significant portion of the adults survived but many of whom suffered uncoordinated movement suggestive of a *js*-like but milder defect (compare the first and second columns in the top row of Figure 6A). JS overproduction driven by tub-Gal4 resulted in a similar but milder phenotype than the act-Gal4 driver (top row, left panel of Figure 6A).

The two CBD-truncated JS proteins, when overproduced, also led to physiological consequences. Overproduction of both proteins had a similar effect as JS-dsRED overproduction when act-Gal4 was employed (top row in Figure 6A). Although overexpression of the two classes of dsRED-tagged JS proteins (wildtype or CBD-deleted) overwhelmingly yield “uncoordinated” adults, a unique wing phenotype was only produced by the latter. Essentially all surviving flies have their wings spread horizontally (Figure 6B). The results from tub-Gal4 driven overexpression were largely consistent with ones from act-Gal4, except that overexpressing JS-dsRED had a much milder effect resulting many normal looking adults (top row in Figure 6A). As all *UAS*-containing constructs were inserted into the same genomic location of 75B via phiC31 integrase mediated insertion, chromosomal position effect on the transgenes must not have been the underlying cause for this phenotypic variation. We suggest that it is more likely caused by the difference in tissue specificities of the two Gal4 drivers.

We hypothesized that JS overexpression might have interfered with the normal function of JS leading to phenotypes similar to JS loss of function. We investigated this potential “dominant negative” nature of JS overproduction by repeating it in a *js* heterozygous background, and expected an enhancement of the negative effect of JS overproduction. Our hypothesis is largely supported by the results. As shown in Figure 6A (compare results in top and bottom rows), adult lethality was driven to essentially 100% with JS-dsRED regardless of which driver was deployed. Even over-expressing JS^⊗CBD-I^-dsRED in a *js*-heterozygous background was sufficient to result in complete lethality. Remarkably, overproduction of JS^⊗CBD-II^-dsRED produced similar phenotypes in the two *js* background: no lethality but all uncoordinated. This suggests that the “spread-wing” phenotype is likely the result of a “gain-of-function”, instead of a “dominant negative” effect from overproducing a defective and possibly mis-localized JS protein. Interestingly, ⊗CBD-II deletes one of the N-linked glycosylation sites (Figure 2A), implying the importance of this modification to JS function. In summary, the above results suggest that overexpression of even partially functional JS proteins disrupts the endogenous JS functions, consistent with that JS overexpression exerts a poisoning effect.

### The genetic determinants of JS localization

To provide further supporting evidence that JS overproduction interfered with the normal JS functions, we studied the localization of dsRED-tagged JS proteins ectopically produced from act-Gal4 in tissues using live fluorescence. We discovered that in most of the tissues where we previously observed JS-mCherry signals (at the endogenous level), JS-dsRED is also present but with a much stronger intensity. Some of the examples are shown in Figure 6C. In particular, the puncta appearance of JS-mCherry molecules changes to a more intense and “sheet” like appearance of JS-dsRED, possibly due to the large amount of JS-dsRED or its mis-localization in the tissues of interest. Interestingly, ectopic JS-dsRED signals were also observed where normal JS-mCherry signals were not discernable, e.g., the secretory cells of the larval salivary glands (Figure 6C), suggestive of protein mis-localization. The localization of the two CBD affected JS proteins are similar with each other but differ mostly from the pattern of the endogenous JS protein. Most prominently, neither is discernable in pericardial cells where endogenous JS is present. In addition, large aggregation of CBD-affected JS-dsRED proteins are present in salivary glands and trachea of the larva. This aggregation is most prominent for the JS^⊗CBD-II^-dsRED protein (Figure 6C). These results are consistent with that the CBD domain is critical for JS localization. They also support our previous propositions that JS^⊗CBD-II^-dsRED is a defective protein, and that overexpression of a normal JS protein is needed to effectively disrupt the function of the endogenous JS.

### Relationship of JS with the previously identified Mucin-D protein

A glycosylated protein named Mucin-D has been extensively characterized by Dr. A. Kramerov and colleagues (e.g., Kramerov et al. 1996; 1997). Several pieces of evidence support the hypothesis that Mucin-D and JS are related, and possibly the same protein. (1) Mucin D was previously purified from various lines of cultured Drosophila cells. It was characterized as a secreted glycoprotein, a character shared with JS. (2) Radio-labelled sugar was used to estimate the molecular size of Mucin-D as being between 70 and 100KD depending on running conditions of SDS-PAGE electrophoresis. Interestingly, Mucin-D molecules sometimes showed two migrating bands around 70 and 100 KD in size, similar to JS’s behavior on an SDS-PAGE. (3) Remarkably, the determined amino acid compositions of Mucin-D match those of JS very well in that the percentages of different residues between the proteins are correlated with very high significance (Spearman’s test, *r_s_* = 0.8135*, p*(2-tailed)=7E-05, Supplemental Materials Table S1).

IgM antibodies against Mucin-D using semi-purified native protein as antigens were previously generated. Unfortunately, none of the ones we tested recognizes JS on Western blots even when we used fly extracts overexpressing JS. Therefore, whether JS and Mucin-D are the same protein required further investigations.

## Discussion

In this study, we characterized lethal mutants with a novel adult phenotype, and identified the affected *js* gene encoding a secreted glycoprotein that bears basic domain features of mammalian Mucins. We also studied the extensive localization of the JS protein in different tissues and developmental stages. Although the exact physiological function of JS remains undetermined, it is likely to be a widely distributed Mucin-like molecules in Drosophila, one of the best models for developmental studies with facile genetics. Future investigations into the consequences of both the loss and gain of JS function promise to further our understandings of Mucins in development and disease.

### JS is a new member of the Mucin-like proteins in Drosophila

Bioinformatic and limited functional studies have led to the identification of many proteins with signatures of mammalian Mucins in Drosophila and other insects (Schwientek et al. 2007; Syed et al. 2008; Dias et al. 2018). Interestingly, JS was not among the previously identified ones, suggestive of the limitation of bioinformatic and genome-wide analyses. In addition to having the most basic features of non-membrane associated Mucin-like proteins (secreted, glycosylated, having an N- and a C-terminal Cysteine-rich domains, and most importantly, having a middle PTS-rich domain), JS has a distribution pattern highly consistent with the general function of Mucin. It is present at the cuticle, digestive, respiratory and reproductive tracts, where the epithelia interact with the environment. In addition, JS is present at internal epithelia in many organs, suggestive of specialized functions, e.g., joints between leg segments, bases at bristles and wing blades, spaces between ommatidia. Therefore, results from our localization studies of JS are consistent with JS serving a similar function as general Mucins in other organisms. In addition, based on a highly correlated amino acid composition between JS and the previously identified Mucin-D protein, we proposed that *js* likely encodes Mucin-D, which Mucin-like properties have been extensively characterized by Kramerov and colleagues. Therefore, JS might represent an excellent example of a general Mucin-like molecule in insects.

### Potential functions of JS

The facts that JS’s presence is widespread early in development and that it persists throughout organogenesis seem inconsistent with the late manifestation of physiological defects in mutant adults. In fact, we argued that protein localization that we presented for JS-mCherry in Figures 3 and 4 is unlikely to reflect the entire JS distribution pattern as we relied entirely on JS-mCherry’s ability to fluoresce under a molecular environment that we have very little knowledge about. One possibility is that defects in *js* mutants are sub-lethal earlier on in development, but rearing conditions under the laboratory settings relax the requirements for JS functions. In other words, more sensitive assays are needed to reveal defects in early *js*-mutants to justify the omni-presence of this protein. For example, the presence of JS in the respiratory and digestive systems might suggest a protective role, the loss of which might not have damaged the luminal surfaces as we confirmed in EM studies. Physical and/or chemical challenges might need to be applied to reveal protective functions that are compromised in the mutants. Moreover, whether the prominent presence of JS in nephrological cells carry functional significance requires further investigation. Alternatively, it might simply reflect that JS is circulating in the hemolymph as excess proteins in the hemolymph are shown to be filtered by the nephrocytes (Weavers et al. 2009; Zhang et al. 2013b).

Interestingly, JS proteins are internal to several different body structures, e.g., at spaces between ommatidia, at the bases of bristles, and at the junctions between leg segments. It is plausible that JS serves the function of a lubricant to facilitate movements of body parts, and the lack of lubrication might have been the underlying cause for breakage of leg joints giving rise to the most distinct phenotypes of *js* adults. Specialized assays would be needed to determine whether lubrication for organ movements is disrupted during earlier development of *js* mutants.

The observation that overexpressing JS or derivatives disrupts the function of the endogenous protein suggests interesting possibilities of JS regulation. First, the presence of excess JS might be sufficient to disrupt the molecular stoichiometry of the mucin barrier. Secondly, the abnormally produced JS might lack functionally important post-translational modifications and/or processing as the acting cellular machineries are likely overwhelmed by the mass production of these secreted molecules. A similar effect on protein folding might exist when JS is being overproduced. In sum, improperly processed and/or mis-folded JS molecules might be incorporated into the epithelial networks where JS is a normal component, thus disrupting these networks.

### The future of JS studies

Our JS study lays the groundwork for future structural and functional studies of Mucin-like molecules in a genetic model. In addition, both loss and gain of function tools are available. (1) On the issue of targeted localization, how does JS achieve its precise distribution? As JS has a well conserved CBD (Jasrapuria et al. 2010; Tetreau et al. 2015), it is highly likely that chitin-binding serves as a primary localization mechanism for JS. In fact, the two CBD-deleted JS-dsRED derivatives show signs of mis-localization (Figure 6C), which is consistent with a critical role of chitin-binding in JS localization. However, the Cys-rich N-terminal domain of mammalian Mucins participates in inter- and intra-molecular crosslinking. Whether JS’s CBD has a similar function and how the JS-chitin and JS-JS modes of interaction are coordinated would be of future interest. How the conserved Cys-rich motif at the C-terminus determines JS junction also awaits further investigations. (2) On the issue of post-translational modifications, how does the extent of JS glycosylation affect its function? The expected O-linked glycosylation sites reside in the PTS-rich domain in the middle of JS. Strategically placed in frame deletions would help identify functionally critical glycosylation positions and scope. In addition, identifying the glycosylase(s) responsible for modifying JS would be of future interest. As most if not all of the enzymes responsible for O-linked glycosylation have been identified in Drosophila (Tran et al. 2012; Zhang et al. 2019), RNAi knockdown combined with a PAGE-based glycosylation assay would yield important insights. (3) On the issue of JS secretion, what are the major cell types responsible for JS secretion? Do these cells carry similar characteristics of mammalian goblet cells that specialized in Mucin secretion?

### Data availability

Strains and plasmids are available upon request. The authors affirm that all data necessary for confirming the conclusions of the article are present within the article, figures, and tables.

## Acknowledgements

We thank Mr. Yuanlong Zhao at the National Cancer Institute, USA for his assistance in the initial mapping of the original *js* mutation. We thank Dr. Andrei Kramerov for kindly providing Mucin-D antibodies. We thank Hongmei Li at the Sun Yat-sen University microscopic core facility for her assistance in EM operations.

## Supplemental Materials

**Figure S1. Signal peptide prediction for JS**

A SignalP-4.1 output predicting the position of the signal peptide in JS protein. The X axis lists the N-terminal residues of JS. The predicted excision position of the signal peptide is marked with the longest vertical red line.

**Figure S2. Amino acid sequence alignment of JS from insects**

Clustal Omega alignment output of JS proteins from selected insect species. “mel”: NP_650538.1 of *Drosophila melanogaster*; “LuciliaCuprina”: XP_023308467.1 of *Lucilia cuprina*; “StomoxysCalcitrans”: XP_013099154.1 of *Stomoxys calcitrans*; “CeratitisCapitata”: XP_012157262.1 of *Ceratitis capitata*; “AedesAegypti”: EAT42245.1 of *Aedes aegypti*; “BombyxMori”: XP_004927149.1 of *Bombyx mori*; “PlutellaXylostella”: KAG7299243.1 of *Plutella xylostella*.

**Figure S3. JS-mCherry fluorescent signals are distinct from auto fluorescence**

Images of the left panel were taken from *js^mCherry^* animals, and shown in triplets. “Green fluorescence” was used to indicate auto-fluorescence. Images on the right were taken from *w^1118^* animals, and shown in triplets. Red signals are the result of auto-fluorescence. Scale bars indicate 100μm.

**Figure S4. JS-EGFP localization in adult eyes**

Images are shown in triplets: GFP fluorescence, brightfield (BF), and the merged image of the two. One of the ocelli is marked with an arrowhead. Scale bars indicate 100μm.

**Movies S1, S2. Eclosion of wildtype and *js* mutant adults**

Two movies recording the eclosion of a wildtype and a *js*-mutant adult.

**Table S1. Amino acid compositions of Mucin-D and JS**

**Table S2. Primer list**

