## supplemental figures and tables for "JiangShi: a widely distributed Mucin-like protein essential for Drosophila development"

### Supplemental Figure Legends

#### Figure S1. Signal peptide prediction for JS

A SignalP-4.1 output predicting the position of the signal peptide in JS protein. The X axis lists the N-terminal residues of JS. The predicted excision position of the signal peptide is marked with the longest vertical red line.

#### Figure S2. Amino acid sequence alignment of JS from insects

Clustal Omega alignment output of JS proteins from selected insect species. “mel”: NP_650538.1 of *Drosophila melanogaster*; “LuciliaCuprina”: XP_023308467.1 of *Lucilia cuprina*; “StomoxysCalcitrans”: XP_013099154.1 of *Stomoxys calcitrans*; “CeratitisCapitata”: XP_012157262.1 of *Ceratitis capitata*; “AedesAegypti”: EAT42245.1 of *Aedes aegypti*; “BombyxMori”: XP_004927149.1 of *Bombyx mori*; “PlutellaXylostella”: KAG7299243.1 of *Plutella xylostella*.

#### Figure S3. JS-mCherry fluorescent signals are distinct from auto fluorescence

Images of the left panel were taken from $js^{mCherry}$ animals, and shown in triplets. “Green fluorescence” was used to indicate auto-fluorescence. Images on the right were taken from $w^{1118}$ animals, and shown in triplets. Red signals are the result of auto-fluorescence. Scale bars indicate 100μm.

**Figure S4. JS-EGFP localization in adult eyes**

Images are shown in triplets: GFP fluorescence, brightfield (BF), and the merged image of the two. One of the ocelli is marked with an arrowhead. Scale bars indicate 100μm.

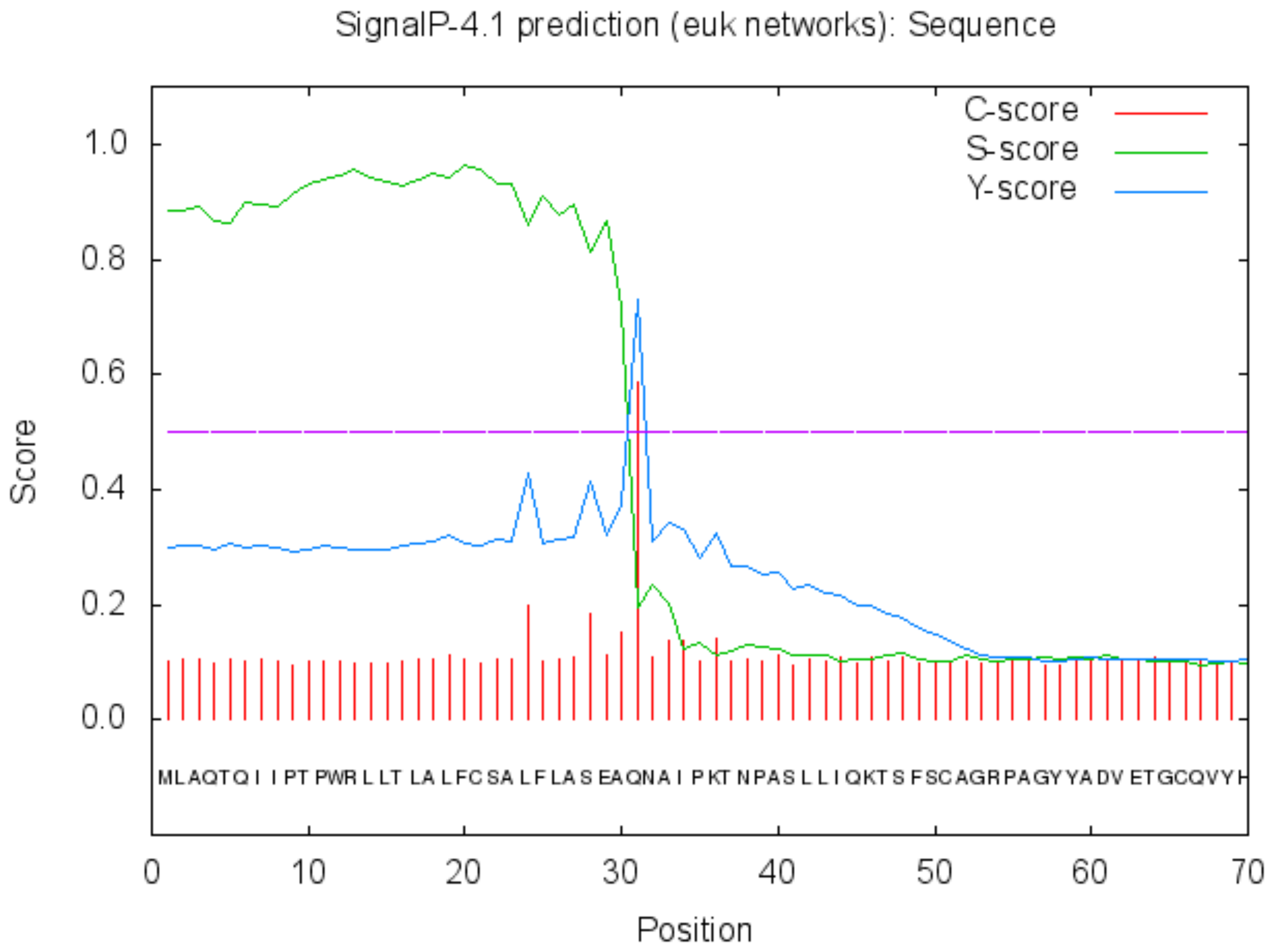
SignalP-4.1 prediction (euk networks): Sequence
C-score
S-score
Y-score
Score
1.0
0.8
0.6
0.4
0.2
0.0
MLAQTQIIPTPWRLLTLALFCSALFLASEAQNAIPKTNPASLLIQKTSFSCAGRPAGYYADVETGCQVYH
0
10
20
30
40
50
60
70
Position

```
mel                      ---------------------------------------------------mlaq-----   4
LuciliaCuprina           ------------------------------------------------------------   0
StomoxysCalcitrans       ------------------------------------------------------------   0
CeratitisCapitata        ------------------------------------------------------------   0
AedesAegypti             -------------------------------------------malpfqshllpti---n   14
BombyxMori               ------------------------------------------------------------   0
PlutellaXylostella       meeltkmmkeiqeelveqkvefqqmernittninnninnkfermeikyaelektvtkqee   60

mel                      -tqiip-----------------tpwr-llt---lalfcs--alflaseaqnaipktnpa   40
LuciliaCuprina           ---mvn-----------------hnwf-latvllwlllc-yhnlcinaqsqnaig----a   34
StomoxysCalcitrans       ---mfn-----------------hnwf-vptvvlsllls-yrnffitaqsqnaig----a   34
CeratitisCapitata        ---mcn-----------------hnwl-twpliiwlltccyflvgidgqsenaig----a   35
AedesAegypti             hnaylaadfrde-----------rrylfvss----vfls-----pqrsqvln-----dgs   49
BombyxMori               -----------------------macerlcvcl--illc---lyciagyaqs-----dtp   27
PlutellaXylostella       rldmiekqlrkknlvffgveeteknyedlesnlvkiind---kmkvscslse-----dvr   112
                                                             :              .

mel                      slliqktsfscagrpagyyadvetgcqvyhmcdglgrqfsytcpnttlfqqrmlicdhwy   100
LuciliaCuprina           hrtfqktsfscagrpsgyyadietgcqvyhmcdglgrqfsyscpnttlfqqrmlicdhwy   94
StomoxysCalcitrans       hrsfqktsfscagrpagyyadvetgcqvyhmcdglgrqfsyscpnttlfqqrmlicdhwy   94
CeratitisCapitata        hrvfektsfscagrpagyyadietgcqvyhmcdglgrqfsyscpnttlfqqrmlicdhwy   95
AedesAegypti             fkfaaktsfscsgraagyyadvetgcqiyhmcdglgrqfsyacpnttlfqqrmlicdhwy   109
BombyxMori               qvkipptsftcrgraagyyadmetgcqvyhmcdglgrrfsyscpkttlfqqrmlvcdhwy   87
PlutellaXylostella       dsripatrfscrgrgsgyyadvetgcqvyhmcdglgrqfsytcpnttlfqqrmlvcdhwy   172
                               * *:* ** :*****:*****:*********:***:**:*********:*****

mel                      mvncskaesnyaanlligqrdkpfvndeenslrtprpdlldrpyapdysgesfrsqykq-   159
LuciliaCuprina           mvncskaesdytanlligqrdkpfvndeenslrtprpdlldrpyapdysgesfrnqyqks   154
StomoxysCalcitrans       mvncskaesdytanlligqrdkpfvneeennlrtprpdlldrpyapdysgesfrsqyqks   154
CeratitisCapitata        mvncsraesdyaanlligqrdkpfvndeenslrtprpdlldrpyapdysgesfrnqykq-   154
AedesAegypti             mvncskaesnyaanlligqrdkpfvtddenelrtprpdlldtpyaagynmds----fkyn   165
BombyxMori               mvncsmaerdydanlligqrdkpfvsdeemsqrtprpdilsvpltskyydglkeaeskfl   147
PlutellaXylostella       mvncsrserdydanlligqrdkpfvsehemqfrtprpdilsvppnsnyydglkeaeskyp   232
                         ***** :* :* *************.:.* . ******:*. *    *         :

mel                      ft-----snqnqirdesvkgaga--------gksdpqisq------trwripppsrtilp   200
LuciliaCuprina           ls-----svlnqiydhtaqkmke---ktlqaqqnqlgtgq------qrwkipppsriilp   200
StomoxysCalcitrans       ls-----svlnqiydtnsykkek---slsssnqptqptgq------qrwkipppsrlilp   200
CeratitisCapitata        lp-----aiqnqiqdtdaqkfqd---klnt--ipqaapgq------qhwkipspsrtilp   198
AedesAegypti             yfktntqapqnaipangkssvktqtvkpkqttseifeesasnhglpihwstryaked---   222
BombyxMori               l------hpdndivgvad--------tisgdednildsakqnyrpptswstrkrr-----   188
PlutellaXylostella       v------hpgnsivgvad--------slssnnngldhgkqqqyrpptswstgtgtrr---   275
                                   * *                                   *

mel                      payepqielpsaqsakpr----ipiitstttttra------ttttrpttttrattttttt   250
LuciliaCuprina           payepqilees----sakpstrtsyytptitt------tt------rkpst-lrptat-t   242
StomoxysCalcitrans       payepqipdntvitttaksttrisyynpsttt------ttttttskpspss-lrptssns   253
CeratitisCapitata        payeaqttddrasqtnsdnv----rfvpktaqptasskqtkttfnrptpss-vsa--stq   251
AedesAegypti             ddkerkd--d-----kqeldtplstaiplste-eaqlqgsktn-er-------rdvkene   266
BombyxMori               p-----------------------------------------------------------   189
PlutellaXylostella       ptsgptr--p-----tpq------------------------------------------   286
```

```
mel                   rrppvtarpkealhnrrpnfqe-------------h---------------dmddlgtsh    282
LuciliaCuprina        knqstkfnnvaalhnrhdehlq-------------l---------------ehddlgtsh    274
StomoxysCalcitrans    rslgtrfngvaalhnrqdehlq-------------y---------------elddlgtsh    285
CeratitisCapitata     vqpinrfnn-aalhqr---gqs-------------n---------------epedlgtsh    279
AedesAegypti          nrttvrvnakekqsnrveeaasnnnqvnvkvpsiryeppftpeeqvkrgsasketlgagf    326
BombyxMori            ---nvqinnid----slagaasq-----epgptirpqpes--------------------    217
PlutellaXylostella    --apsgidaqd----nlagaasq-----kptptyqtlpdf--------------------    315
                                           .

mel                   stryntsadfnsaesplretkqsst-----------------------------------    307
LuciliaCuprina        stryntsadfnsaedrfaktpqkatt-prkstynsnfnnaft------------------    315
StomoxysCalcitrans    stryntsadfnsnedrnqknrgknaytttkssfg--lttlrt------------------    325
CeratitisCapitata     stryntsadfnsaeqpnsktasvsnknkisssfgivppatastt------ttttrpvtpt    333
AedesAegypti          -kpfesilnfnkkgnrkd--ydfsnlfgrnesirntppaqtptpastttprtiqssttrf    383
BombyxMori            -------rdtslk-------------n------kgnqperelmppllp--stteae----    245
PlutellaXylostella    -------sdfnqn-------------tgvpnsrpvspptqdlrppeap--ttteqp----    349
                              : .

mel                   -----------------------------kltkfikppskiyeppfvypiynleesqtq-    337
LuciliaCuprina        ----takptt--------------ttkkttlnpkikvpskiyepplly piynmddnta--    355
StomoxysCalcitrans    ----ttstst--------------sttkatanpnikipskiyeppvlfpiynledssttt    367
CeratitisCapitata     asskltttsst-------------ssaaittnlpikipskvyeppyvypiynenetvst-    378
AedesAegypti          fsssssttpsrsttrfsstsrqtgfvtstgttqkpisiivsdlqppqlnpaqfsprtgp--    441
BombyxMori            ----------d---------------datttqssvyqfikr----------fdpnspd--    268
PlutellaXylostella    ----------sseeydld------irmknssedpvfqfikr----------fdpnspd--    381
                                                   .    :    .               .

mel                   -naavattlrtstaapfspvpsrke-vstttprplsrpttlagvpfssathpttvttv--    393
LuciliaCuprina        -tttmrptfrastaspfvtrvsat-ts-ttttkaptrpttvigkpftrssfstst----k    408
StomoxysCalcitrans    ttttvrplfrastaapfmtrqttt-ttrptttrppsrattmlgkpftrsssitapt---t    423
CeratitisCapitata     -----qlplrassatpftvsptltrltsttpvpppsrpttlagkpfaapssgittkapnf    433
AedesAegypti          ---afrpavt--taaprstftata-ttttttpr----pltlsdpr---q-snrapttilf    487
BombyxMori            ---siktait------qseiidl----nkhlpe----gqvssegqvsse-dertpkk---    307
PlutellaXylostella    ---slkttmt------aseilni----nqqlpq----gqd-----vste-eertprr---    415
                              .                                            :

mel                   ----g-----------vpprsdnrtpapaqsfrlatptstaa------------------    420
LuciliaCuprina        v---gtnsnnidrfdvsklskeirtpapaqgfkpptptppvknsrtsqnstynkp-----    460
StomoxysCalcitrans    t---tqsgrnlnrfdadklskelrtpapaqgfkiptprptvsafnts-------------    467
CeratitisCapitata     g---grsqnnliqnsfthnskdvrtptpaqgfkpptprplistttsatkstesspfdysi    490
AedesAegypti          nppaataglfenrfalqrppvtskvpvpsrdilppf--qnvanfdkiknrfv--------    537
BombyxMori            ------nksfgnklndfk-----------tdndkkf--regtrfdsvntk----------    338
PlutellaXylostella    ------qknrgnnfntlq-----------tekrkgn--teqnrfntintk----------    446

mel                   -----------------------------------------ppsrpaqlpfndllppfvd    439
LuciliaCuprina        ----------------------------------------drgsvkheqpskqllpplad    480
StomoxysCalcitrans    ------------------------------------------ksiaesvpskhllpplvd    485
CeratitisCapitata     pttnrlhlptpiintasstn----------------tgtaaatgsdatppskellppyqe    534
AedesAegypti          --seaiftrqpitnepssgpiksgidklqgdvqiqvrndnvprpfsvptpandllppkke    595
BombyxMori            --------pnptennt---------------------egdsgvkskvpkpeqvllppkkd    369
PlutellaXylostella    --------ptaptplpdtnnfspgn------------dnsnqrstglsepdkhllppksd    486
                                                                       * : ****  :
```

```
mel                 fvph-diattqgppiyyewk--vpsngleppkldppigvdgreypettgdygvtskqdvf    496
LuciliaCuprina      fith-dvattqgppiyyewk--vpsdgleppkldspigvdgreypetvidynsistnngf    537
StomoxysCalcitrans  fvqh-dvattqgppiyyewk--vpsdgleppkmevpigvdgrqypetvldynsisasgtf    542
CeratitisCapitata   fvqh-diattqgppiyyewk--vpsngleppkldppigvdgreypepsfdyntisgaseg    591
AedesAegypti        yvfy-ddattkgppiyyqwkwsipsfgleppldapltdeek--llt--------------    638
BombyxMori          ysqspldsttmgppiyyewkwavpafgllppkegnitnsta--lp---------------    412
PlutellaXylostella  yavi--psttvgppvyyewkwavpafdlelplagnstdnst--aae--------------    528
                    :      :** ***:**:**  :*: .*  *      .

mel                 ntrlndigshqkkpvqitsplqqsstshrlaisrsikpkeeqesaq--------------    542
LuciliaCuprina      tsi-ssfgidqkrsnntsksenktivpsrspisrsiketgqpkgvnetkak-----tsnt    591
StomoxysCalcitrans  spn-phfgidqkkevdsr-------ptlrpptsrsvketaqpngvstrrka-----dkpt    589
CeratitisCapitata   knt-qt---aksnssptdkvasdksqtartpisrsiketgqpnsvhrvvnvestkatstt    647
AedesAegypti        --q------lnpasqdspdsnrehnqtarsiss----etgnpgenp-----------sts    675
BombyxMori          -----------tkspk--tgkspfs-tarrtti----kaeda------------------    436
PlutellaXylostella  --v------pqgrsns--qgnrpfs-nwsrptt------eep------------------    553
                                                            :

mel                 ------rrsdvvasstdishlrkqflipeyafpletigr--tgygpgagaaaggsgssng    594
LuciliaCuprina      sptrastttqkssnpgdifqirkelsvpeyafplenvgr--tgyl-------------dt    636
StomoxysCalcitrans  k--atgtaptssnepedifqiskdlsvpeyafplenvgr--tgyl-------------ds    632
CeratitisCapitata   tpvatdrsdtisaptsditqlrrdysvpeylfplesagr--tgyl-------------st    692
AedesAegypti        ntiaeankrkiqlpthnylelrkqlaipdftfplesagtaqsvye-------------qe    722
BombyxMori          ----tpkprnve-------ynissyyvpdyvfpldgp---hpgye-------------sd    469
PlutellaXylostella  ----dlnpsnve-------ynissyfvpdyqfpldkp---hpgye-------------de    586
                                           .  :*:: ***:        *              .

mel                 dlynsfqlkipeqra-------kwfgenpkcpechpsfvlpgtcepclrr--    637
LuciliaCuprina      dvynsfqlkiperra-dtdehihwygenpkcpechpsyvrpgtcepcirr--    685
StomoxysCalcitrans  dvynsfqlkiperra-dtdehlhwfgenpkcpechpsyvipgtcepclrr--    681
CeratitisCapitata   dsynsfrlkipehgeeeiedhrhwfgenpkcpechpsfvvpgtcepcirr--    742
AedesAegypti        gavnsfqikipayar-sdddsrawygenakcpqchpafmrpgscepcikirr    773
BombyxMori          daqtsfqvqvsr-------pgrssygenpkcpqchpaylkpgscepcivkr-    513
PlutellaXylostella  naetsfqvkvgr-------pgr-sfgenpncphchpaylnpgtcepcivkr-    629
                    .  .**::::              :*** :**.***::: **:****:
```

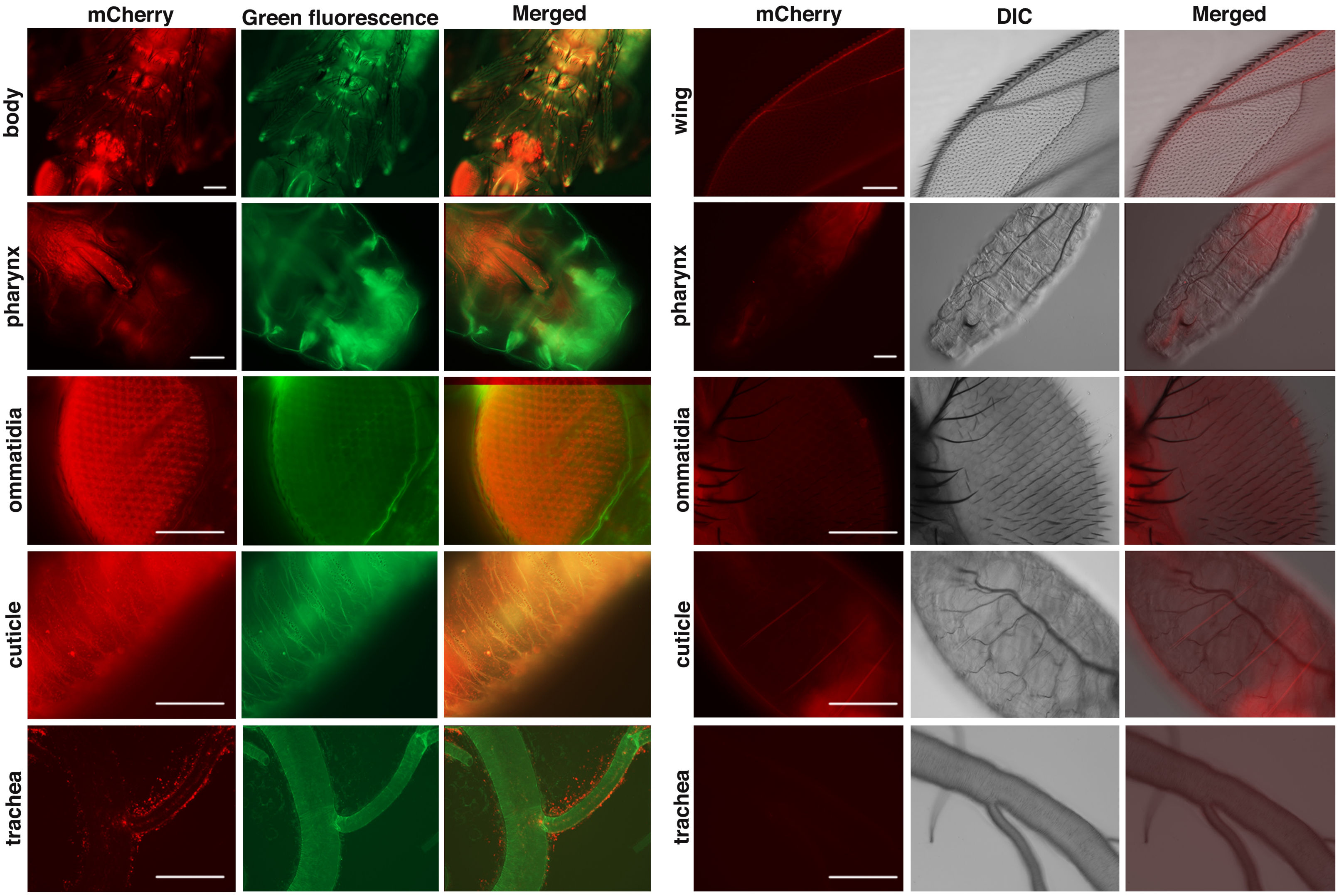
mCherry
Green fluorescence
Merged
mCherry
DIC
Merged
body
pharynx
ommatidia
cuticle
trachea
wing
pharynx
ommatidia
cuticle
trachea

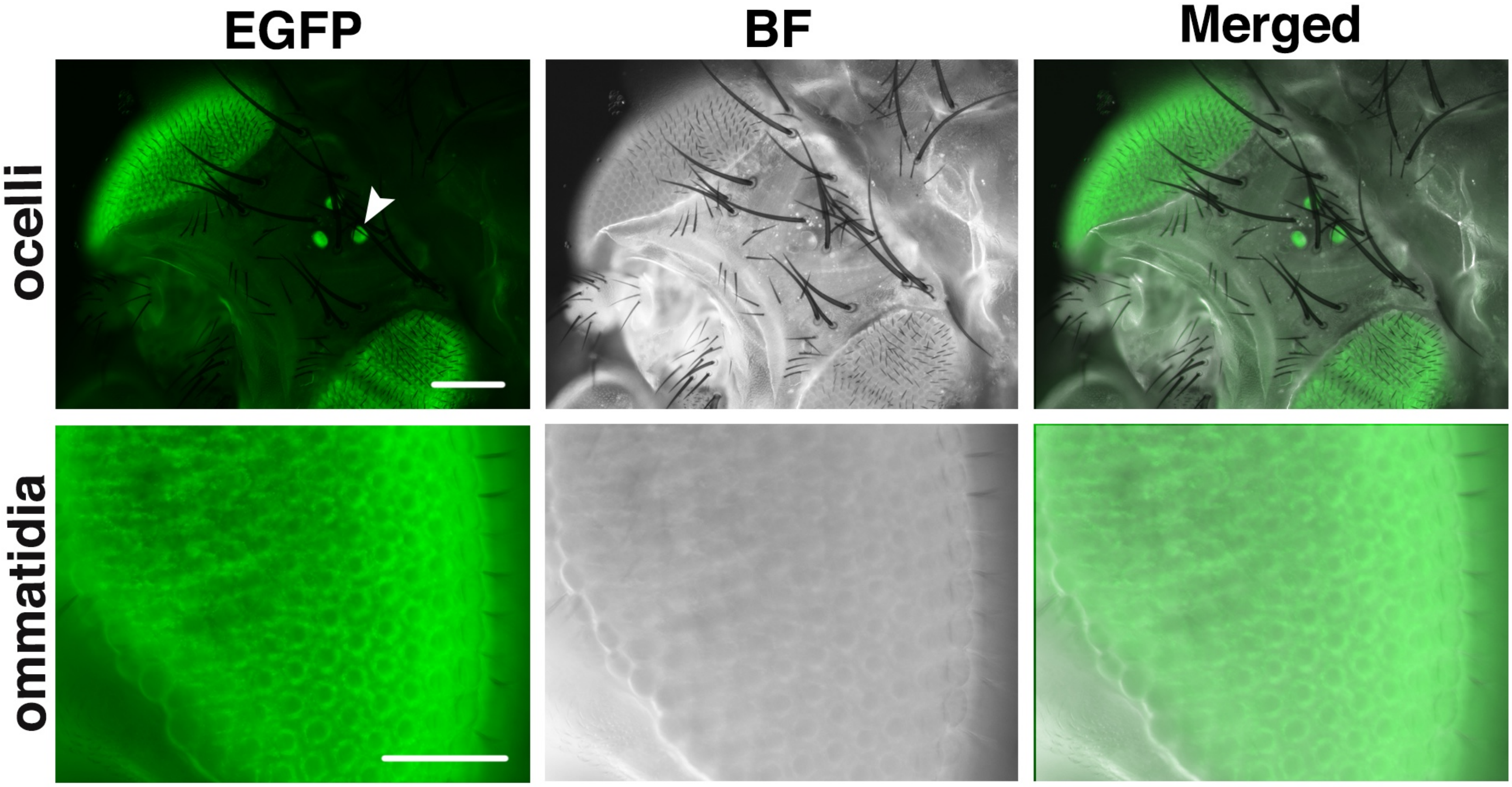
EGFP
BF
Merged
ocelli
ommatidia

Table S1. Amino acid compositions of Mucin-D and JS[4]

| Amino acid composition[1] | Mucin-D[2] | JS[3] |
|---|---|---|
| Asp+Asn | 7.5 | 8.3 |
| Thr | 11.5 | 11.4 |
| Ser | 10.8 | 9.5 |
| Glu+Gln | 10.5 | 10.1 |
| Pro | 7.3 | 11.6 |
| Gly | 15 | 5.1 |
| Ala | 8.4 | 7.1 |
| Cys | 1.5 | 1.7 |
| Val | 4 | 4 |
| Met | 1.1 | 0.7 |
| Ile | 2.2 | 4.3 |
| Leu | 5.5 | 6 |
| Tyr | 1.4 | 3.5 |
| Phe | 3.5 | 3.6 |
| Lys | 4.6 | 4.3 |
| His | 0.9 | 1.7 |
| Arg | 3.3 | 7 |

[1]: Three letter abbreviations of analyzed residues
[2]: The amino acid composition of Mucin-D was taken from Kramerov et al. (1996) as "number of residues per 100 amino acids".
[3]: Calculated amino acid composition of JS protein with the predicted signal peptide residues not included in the calculation.
[4]: A spearman correlation test yields $r_s$ = 0.8135, *p*(2-tailed)=7E-05.

Table S2. Primer list

| **Primer name** | **Sequence** |
|---|---|
| **Construct of pUAST-*js* cDNA** | |
| 14880cDNA FP | ATGTTGGCACAAACGCAAATTATCC |
| 14880cDNA RP | CTATCTGCGCAGGCAGGGCT |
| **Pet28a-JS antigen** | |
| pet28a-EcoRI-14880cDNA FP | ATCTATGAATTCATGTTGGCACAAACGC |
| pet28a-HindIII-14880cDNA RP | ATCTATAAAGCTTCTATCTGCGCAGGCAGG |
| **pUAST-*js* cDNA-dsRED** | |
| 14880-cDNA-dsRED YJ FP | GTTCTGCCGGGCACCTGCGAGCCCTGCCTGCGCAGAGGC GGAGGGGCCTCCTCCGAGAACGTCATCAC |
| 14880cDNA YJ RP | TCTGCGCAGGCAGGGCTCGCAGGTG |
| **N-terminal gRNA primer** | |
| ATG -gRNA-FP | GTCGAGAACAATCGTCGAAATGT |
| ATG-gRNA-RP | AAACACATTTCGACGATTGTTCT |
| **C-terminal gRNA primer** | |
| TAG-gRNA-FP | GTCGGAATCAGGACTATCTGCGC |
| TAG-gRNA-RP | AAACGCGCAGATAGTCCTGATTC |
| ***$Js^{3\text{-}1}$* insert fragment PCR** | |
| 14880-$w^{+}$-FP-2 | ATCCGATGTCGACTCCAACC |
| 14880-exon2-RP-1 | CGAATTGCGAACCGTCAGAT |

**Sequencing results**

(1) CRSPR/Cas9 induced *js* mutations

In the sequences below, the start codon **ATG** is in bold.

TGCCACAGAGAACAATCGTCGAA**ATG**TTGGCACAAACGCAAA *wt*<br>
TGCCACAGAGAACAATCGTC------------------GCACAAACGCAAA $js^{cas-1}$

TGCCACAGAGAACAATCGTCGAA**ATG**TTGGCACAAACGCAAA *wt*<br>
TGCCACAGAGAACAACAATCG------**TG**TTGGCACAAACGCAAA $js^{cas-2}$

In the sequences below, the stop codon **TAG** is in bold.

CACCTGCGAGCCCTGCCTGCGCAGA**TAG**TCCTGATTCCAT *wt*<br>
CACCTGCGAGCCCTGCCTGC---------------**G**TCCTGATTCCAT $js^{c-term}$

(2) *P* element in $js^{3-1}$ (imprecise excision)

**Sequences in bold are from the 5'UTR of CG14880.** <u>Sequence underlined is the 5' end of P element</u>. *Sequences in italic are part of the origin of the yeast 2micron plasmid*.

**GACTAGTGACATGATGTGCTAAAGTGTTGATGGCATGTC**<u>CATGATGAAATAACATAA</u><br>
<u>GGTGGTCCCGTCGATAGCCGAAGCTTACCGAAGTATACACTTAAATTCAGTGCACGT</u><br>
<u>TTGCTTGTTGAGAGGAAAGGTTGTGTGCGGACGAATTTTTTTTTGAAAACATTAACC</u><br>
<u>CTTACGTGGAATAAAAAAAAATGAAATATTGCAAATTTTGCTGCAAAGCTGTGACTGG</u><br>
<u>AGTAAAATTAATTCACGTGCCGAAGTGTGCTATTAAGAGAAAATTGTGGGAGCAGAG</u><br>
<u>CCTTGGGTGCAGCCTTGGTGAAAACTCCCAAATTTGTGATACCCACTTTAATGATTC</u><br>
<u>GCAGTGGAAGGCTGCACCTGCAAAAGGTCAGACATTTAAAAGGAGGCGACTCAAC</u><br>
<u>GCAGATGCCGTACCTAGTAAAGTGATAGAGCCTGAACCAGAAAAGATAAAAGAAGG</u><br>
<u>CTATACCAGTGGGAGTACACAAACAGAGTAAGTTTGAATAGTAAAAAAAATCATTTAT</u><br>
<u>GTAAACAATAACGTGACTGTGCGTTAGGTCCTGTTCATTGTTTAATGAAAATAAGAGC</u><br>
<u>TTGAGGGAAAAAATTCGTACTTTGGAGTACGAAATGCGTCGTTTAGAGCAGCAGCC</u><br>
<u>GAA</u>*TTCCACGGACTATAGACTATACTAGTATACTCCGTCTACTGTACGATACACTTCC*<br>
*GCTCAGGTCCTTGTCCTTTAACGAGGCCTTACCACTCTTTTGTTACTCTATTGATCCA*<br>
*GCTCAGCAAAGGCAGTGTGATCTAAGATTCTATCTTCGCGATGTAGTAAAACTAGCT*<br>
*AGACCGAGAAAGAGACTAGAAATGCAAAAGGCACTTCTACAATGGCTGCCATCATTA*<br>
*TTATCCGATGTGACGCTGCAGCTTCTCAATGATATTCGAATACGCTTTGAGGAGATAC*<br>
*AGCCTAATATCCGACAAACTGTTTACAGATTTACGATCGTACTGGTACCCATCATGAT*<br>
*TTTGACATCGACCTGGGAGTTTCCCTGAAACAGATAGTATAATTGACTGTATAATAATA*<br>
*TATAGTCTAGCGCTTACCGAGACATGATTGTATTTCGTCTGGAAAACTATGCATCATG*<br>
*CATAGGTATCTTGG*

(3) *js* knock-in alleles

In the sequences below, small letters represent genomic sequences from *js*, LARGE LETTERS REPRESENT CODING SEQUENCES FOR FLUORESCENT PROTEINS.

*js*$^{mcherry}$

ccacccctttgtgtaccccatctacaatctagaagagtcccagacccagaacgcggcagtcgccaccacactgcgcacat
caacggcggctcctttcagtcctgttccaagtcgcaaggaggtcagcaccacaactccacgaccactgagtcgaccgac
gaccttggccggtgtacccttctcctctgccacacacccaactacggttacgacggtcggagtaccgccacgcagcgata
atcggactccagctccggcccagagcttccgcctggccacgcccaccagcactgcagcaccaccatcgcgcccggca
cagttgcccttcaacgatttactgccgccgttcgttgactttgtgccccacgatatagccaccacccaaggaccgcctatctat
tacgaatggaaggtgccctcgaacggtcttgagcctcccaaattagacccacccattggtgtggatggacgtgagtatccc
gagaccactggagactacggggtcaccagcaagcaggatgtattcaacacccgactaaacgatattggaagccatcaa
aagaagccagtccagataacctcgccgctccaacagtcgagcacaagccatcgtttggccatctcgagatccattaagc
ccaaagaggagcaggaatcggcacagcggcgatccgatgtggtggccagctccacggatataagccatctgcgcaag
caattcctcattccggagtacgccttcccgctggaaaccattgggcgcacgggttatggtcctggtgcaggagcagcggct
gggggctccggctccagcaacggcgatctgtataactcgttccagctgaagatccccgagcagcgcgctaagtggttcgg
agagaaccccaagtgcccggagtgccatccttcgttcgttctgccgggcacctgcgagccctgcctgcgcagaGGCG
GAGGGGTGAGCAAGGGCGAGGAGGATAACATGGCCATCATCAAGGAGTTCATGCG
CTTCAAGGTGCACATGGAGGGCTCCGTGAACGGCCACGAGTTCGAGATCGAGGG
CGAGGGCGAGGGCCGCCCCTACGAGGGCACCCAGACCGCCAAGCTGAAGGTGA
CCAAGGGTGGCCCCCTGCCCTTCGCCTGGGACATCCTGTCCCCTCAGTTCATGTA
CGGCTCCAAGGCCTACGTGAAGCACCCCGCCGACATCCCCGACTACTTGAAGCTG
TCCTTCCCCGAGGGCTTCAAGTGGGAGCGCGTGATGAACTTCGAGGACGGCGGC
GTGGTGACCGTGACCCAGGACTCCTCCTTGCAGGACGGCGAGTTCATCTACAAGG
TGAAGCTGCGCGGCACCAACTTCCCCTCCGACGGCCCCGTAATGCAGAAGAAGAC
CATGGGCTGGGAGGCCTCCTCCGAGCGGATGTACCCCGAGGACGGCGCCCTGAA
GGGCGAGATCAAGCAGAGGCTGAAGCTGAAGGACGGCGGCCACTACGACGCTGA
GGTCAAGACCACCTACAAGGCCAAGAAGCCCGTGCAGCTGCCCGGCGCCTACAAC
GTCAACATCAAGTTGGACATCACCTCCCACAACGAGGACTACACCATCGTGGAACA
GTACGAACGCGCCGAGGGCCGCCACTCCACCGGCGGCATGGACGAGCTGTACAA
GGGGCGCGCCTAGtcctgattccatatccattcccgtctccgtccgtcccatcgcccattatccgcatccgcatctgc
tactagccctagtattagcttccgctgagaagtcccggagtgaagcatccacgcatttattgtatatccgcagattaattattaa
actacacaaatatgtttgtgaatgatttatttcgcctgtggtcgtgggtcatttgtgtaattttatatgcaaagaatatcttttatgagt
taggctaagatatttcatggggtaacctaaaaaatttgattattaactcttgatagccaatttttcacagttgttaaaattgagac
accactgaaactttcgaactcgaaatttgggcagtcacaccgtaatgatatttctgtctttatggtttttagatcaatttttcggttttt
tcaaaatttattcagatcagtaaaatacagtattatgggggactgcgcagatgcaaaattagttagtcgagttacggtgcca
ccaataccgaaacttggcgacatgcacatgtattgggtgtggcgaatgctgattaaatcatatttggatgctataggtctttgg
tctggaaatcacccaaaggaggtaatatatatactgttaataaatatggtggttaaatctgcacaattcgttattttaagagcg
cccatacaaaatttgtgctgctgagcacgttggaaatatgggttgtaagaagggaatacgatgaaatgagcagcaggtcc
attttcgaagatctcgaggaacgatatgctatcactaacaatgacaaacgaggtcagttggcaatggatagatttatttaaa
atcaataagatttgtttttattagactctatgtacaacttttgattgctgcctggattactactgtagtgtaataaatctcaaacttca
atatataattatatatatttataaatttggtgaaaattgtcgatctcggacggttggatattatattcatttaaaattgtcatggtaa

gcatgcataaaattgtacaaaacacatcacactagcttaaatctactttcaatgaaattatattttaaatatcaaccgaagtcc
cgggaaatatacc

*js*$^{egfp}$

ccacccttgtgtaccccatctacaatctagaagagtcccagacccagaacgcggcagtcgccaccacactgcgcacat
caacggcggctcctttcagtcctgttccaagtcgcaaggaggtcagcaccacaactccacgaccactgagtcgaccgac
gaccttggccggtgtacccttctcctctgccacacacccaactacggttacgacggtcggagtaccgccacgcagcgata
atcggactccagctccggcccagagcttccgcctggccacgcccaccagcactgcagcaccaccatcgcgcccggca
cagttgcccttcaacgatttactgccgccgttcgttgactttgtgccccacgatatagccaccacccaaggaccgcctatctat
tacgaatggaaggtgccctcgaacggtcttgagcctcccaaattagacccacccattggtgtggatggacgtgagtatccc
gagaccactggagactacggggtcaccagcaagcaggatgtattcaacacccgactaaacgatattggaagccatcaa
aagaagccagtccagataacctcgccgctccaacagtcgagcacaagccatcgtttggccatctcgagatccattaagc
ccaaagaggagcaggaatcggcacagcggcgatccgatgtggtggccagctccacggatataagccatctgcgcaag
caattcctcattccggagtacgccttcccgctggaaaccattgggcgcacgggttatggtcctggtgcaggagcagcggct
gggggctccggctccagcaacggcgatctgtataactcgttccagctgaagatccccgagcagcgcgctaagtggttcgg
agagaaccccaagtgcccggagtgccatccttcgttcgttctgccgggcacctgcgagccctgcctgcgcagaGGGC
GCGCCAGCAAGGGCGAGGAGCTGTTCACCGGGGTGGTGCCCATCCTGGTCGAGC
TGGACGGCGACGTAAACGGCCACAAGTTCAGCGTGTCCGGCGAGGGCGAGGGCG
ATGCCACCTACGGCAAGCTGACCCTGAAGTTCATCTGCACCACCGGCAAGCTGCC
CGTGCCCTGGCCCACCCTCGTGACCACCCTGACCTACGGCGTGCAGTGCTTCAGC
CGCTACCCCGACCACATGAAGCAGCACGACTTCTTCAAGTCCGCCATGCCCGAAG
GCTACGTCCAGGAGCGCACCATCTTCTTCAAGGACGACGGCAACTACAAGACCCG
CGCCGAGGTGAAGTTCGAGGGCGACACCCTGGTGAACCGCATCGAGCTGAAGGG
CATCGACTTCAAGGAGGACGGCAACATCCTGGGGCACAAGCTGGAGTACAACTAC
AACAGCCACAACGTCTATATCATGGCCGACAAGCAGAAGAACGGCATCAAGGTGAA
CTTCAAGATCCGCCACAACATCGAGGACGGCAGCGTGCAGCTCGCCGACCACTAC
CAGCAGAACACCCCCATCGGCGACGGCCCCGTGCTGCTGCCCGACAACCACTAC
CTGAGCACCCAGTCCGCCCTGAGCAAAGACCCCAACGAGAAGCGCGATCACATGG
TCCTGCTGGAGTTCGTGACCGCCGCCGGGATCACTCTCGGCATGGACGAGCTGTA
CAAGTAGtcctgatttcatatccattcccgtctccgtccgtcccatcgcccattatccgcatccgcatctgctactagcccta
gtattagcttccgctgagaagtcccggagtgaagcatccacgcatttattgtatatccgcagattaattattaaactacacaa
atatgtttgtgaatgatttatttcgcctgtggtcgtgggtcatttgtgtaattttatatgcaaagaatatcttttatgagttaggctaag
atatttcatggggtaacctaaaaaatttgattattaactcttgatagccaatttttcacagttgttaaaattgagacaccactgaa
actttcgaactcgaaatttgggcagtcacaccgtaatgatatttctgtctttatggtttttagatcaatttttcggtttttcaaaatttat
tcagatcagtaaaatacagtattatgggggactgcgcagatgcaaaattagttagtcgagttacggtgccaccaataccga
aacttggcgacatgcacatgtattgggtgtggcgaatgctgattaaatcatatttggatgctataggtctttggtctggaaatca
cccaaaggaggtaatatatatactgttaataaatatggtggttaaatctgcacaattcgttattttaagagcgcccatacaaa
atttgtgctgctgagcacgttggaaatatgggttgtaagaagggaatacgatgaaatgagcagcaggtccattttcgaagat
ctcgaggaacgatatgctatcactaacaatgacaaacgaggtcagttggcaatggatagatttatttaaaatcaataagatt
tgtttttattagactctatgtacaacttttgattgctgcctggattactactgtagtgtaataaatctcaaacttcaatatataattata
tatatttataaatttggtgaaaattgtcgatctcggacggttggatattatattcatttaaaatttgtcatggtaagcatgcataaa
attgtacaaaacacatcacactagcttaaatctactttcaatgaaattatattttaaatatcaaccgaagtcccgggaaatata
cc

*js*$^{dsRED}$

ccacccttgtgtaccccatctacaatctagaagagtcccagacccagaacgcggcagtcgccaccacactgcgcacat
caacggcggctcctttcagtcctgttccaagtcgcaaggaggtcagcaccacaactccacgaccactgagtcgaccgac
gaccttggccggtgtacccttctcctctgccacacacccaactacggttacgacggtcggagtaccgccacgcagcgata
atcggactccagctccggcccagagcttccgcctggccacgcccaccagcactgcagcaccaccatcgcgcccggca
cagttgcccttcaacgatttactgccgccgttcgttgactttgtgccccacgatatagccaccacccaaggaccgcctatctat
tacgaatggaaggtgccctcgaacggtcttgagcctcccaaattagacccacccattggtgtggatggacgtgagtatccc
gagaccactggagactacggggtcaccagcaagcaggatgtattcaacacccgactaaacgatattggaagccatcaa
aagaagccagtccagataacctcgccgctccaacagtcgagcacaagccatcgtttggccatctcgagatccattaagc
ccaaagaggagcaggaatcggcacagcggcgatccgatgtggtggccagctccacggatataagccatctgcgcaag
caattcctcattccggagtacgccttcccgctggaaaccattgggcgcacgggttatggtcctggtgcaggagcagcggct
gggggctccggctccagcaacggcgatctgtataactcgttccagctgaagatccccgagcagcgcgctaagtggttcgg
agagaaccccaagtgcccggagtgccatccttcgttcgttctgccgggcacctgcgagccctgcctgcgcagaATGGC
CTCCTCCGAGAACGTCATCACCGAGTTCATGCGCTTCAAGGTGCGCATGGAGGGC
ACCGTGAACGGCCACGAGTTCGAGATCGAGGGCGAGGGCGAGGGCCGCCCCTAC
GAGGGCCACAACACCGTGAAGCTGAAGGTGACCAAGGGCGGCCCCCTGCCCTTC
GCCTGGGACATCCTGTCCCCCCAGTTCCAGTACGGCTCCAAGGTGTACGTGAAGC
ACCCCGCCGACATCCCCGACTACAAGAAGCTGTCCTTCCCCGAGGGCTTCAAGTG
GGAGCGCGTGATGAACTTCGAGGACGGCGGCGTGGCGACCGTGACCCAGGACTC
CTCCCTGCAGGACGGCTGCTTCATCTACAAGGTGAAGTTCATCGGCGTGAACTTCC
CCTCCGACGGCCCCGTGATGCAGAAGAAGACCATGGGCTGGGAGGCCTCCACCG
AGCGCCTGTACCCCCGCGACGGCGTGCTGAAGGGCGAGACCCACAAGGCCCTGA
AGCTGAAGGACGGCGGCCACTACCTGGTGGAGTTCAAGTCCATCTACATGGCCAA
GAAGCCCGTGCAGCTGCCCGGCTACTACTACGTGGACGCCAAGCTGGACATCACC
TCCCACAACGAGGACTACACCATCGTGGAGCAGTACGAGCGCACCGAGGGCCGC
CACCACCTGTTCCTGTAGtcctgattccatatccattcccgtctccgtccgtcccatcgcccattatccgcatccgc
atctgctactagccctagtattagcttccgctgagaagtcccggagtgaagcatccacgcatttattgtatatccgcagattaa
ttattaaactacacaaatatgtttgtgaatgatttatttcgcctgtggtcgtgggtcatttgtgtaattttatatgcaaagaatatctttt
atgagttaggctaagatatttcatggggtaacctaaaaaatttgattattaactcttgatagccaatttttcacagttgttaaaatt
gagacaccactgaaactttcgaactcgaaatttgggcagtcacaccgtaatgatatttctgtctttatggtttttagatcaattttt
cggtttttcaaaatttattcagatcagtaaaatacagtattatgggggactgcgcagatgcaaaattagttagtcgagttacg
gtgccaccaataccgaaacttggcgacatgcacatgtattgggtgtggcgaatgctgattaaatcatatttggatgctatagg
tctttggtctggaaatcacccaaaggaggtaatatatatactgttaataaatatggtggttaaatctgcacaattcgttattttaa
gagcgcccatacaaaatttgtgctgctgagcacgttggaaatatgggttgtaagaagggaatacgatgaaatgagcagc
aggtccattttcgaagatctcgaggaacgatatgctatcactaacaatgacaaacgaggtcagttggcaatggatagattt
atttaaaatcaataagatttgtttttattagactctatgtacaacttttgattgctgcctggattactactgtagtgtaataaatctca
aacttcaatatataattatatatatttataaatttggtgaaaattgtcgatctcggacggttggatattatattcatttaaaatttgtc
atggtaagcatgcataaaattgtacaaaacacatcacactagcttaaatctactttcaatgaaattatattttaaatatcaacc
gaagtcccgggaaatatacc

(4) *js* overexpression constructs

*UAS-js*

In the sequences below, small letters represent pUAST sequence. LARGE LETTERS REPRESENT *JS* CDNA SEQUENCE.

acaggcggcagctgacagctaaacaatctgcagtaaagtgcaagttaaagtgaatcaattaaaagtaaccagcaacca
agtaaatcaactgcaactactgaaatctgccaagaagtaattattgaatacaagaagagaactctgaatagggaattggg
aattcgttaacagatctgATGTTGGCACAAACGCAAATTATCCCAACGCCTTGGCGACTTTTG
ACCCTGGCTTTATTCTGCTCAGCTTTATTTTTGGCCAGTGAGGCTCAAAATGCCATC
CCAAAAACGAATCCTGCCTCGCTGCTGATACAGAAGACATCCTTCTCCTGCGCCGG
ACGTCCAGCCGGATATTATGCGGATGTGGAGACGGGCTGCCAGGTGTACCACATGT
GCGATGGCCTGGGTCGCCAGTTCAGCTACACCTGCCCAAACACGACACTTTTCCA
GCAGCGAATGCTTATCTGCGACCACTGGTACATGGTGAACTGCTCCAAGGCGGAG
AGCAACTATGCTGCCAATCTCCTAATTGGTCAGCGGGACAAGCCCTTCGTAAACGA
CGAGGAAAACAGCTTGCGCACTCCAAGACCCGATCTTCTGGATCGTCCTTATGCGC
CCGACTATTCCGGCGAGTCCTTCAGAAGCCAATATAAGCAGTTTACTTCCAACCAGA
ATCAGATACGTGATGAGTCCGTCAAAGGAGCCGGTGCGGGTAAATCGGATCCCCA
GATATCGCAGACGCGTTGGCGCATTCCACCACCCAGCCGGACGATCCTTCCACCG
GCCTATGAACCGCAAATCGAGCTGCCCAGTGCCCAATCGGCCAAGCCCAGGATAC
CCATTATCACTAGCACCACCACAACCACTCGAGCGACCACCACCACCCGACCAACA
ACCACAACTCGTGCCACCACCACAACAACAACCACTCGAAGACCTCCGGTAACCG
CCAGGCCGAAGGAGGCCTTGCACAACAGGCGGCCAAATTTCCAGGAGCATGATAT
GGATGACTTGGGCACCAGCCACAGCACACGGTACAACACCTCGGCAGACTTCAAC
TCGGCGGAGTCGCCGCTCCGTGAAACGAAACAGTCCAGCACCAAGCTGACCAAGT
TCATCAAGCCGCCGTCCAAGATCTATGAGCCACCCTTTGTGTACCCCATCTACAATC
TAGAAGAGTCCCAGACCCAGAACGCGGCAGTCGCCACCACACTGCGCACATCAAC
GGCGGCTCCTTTCAGTCCTGTTCCAAGTCGCAAGGAGGTCAGCACCACAACTCCA
CGACCACTGAGTCGACCGACGACCTTGGCCGGTGTACCCTTCTCCTCTGCCACAC
ACCCAACTACGGTTACGACGGTCGGAGTACCGCCACGCAGCGATAATCGGACTCC
AGCTCCGGCCCAGAGCTTCCGCCTGGCCACGCCCACCAGCACTGCAGCACCACC
ATCGCGCCCGGCACAGTTGCCCTTCAACGATTTACTGCCGCCGTTCGTTGACTTTG
TGCCCCACGATATAGCCACCACCCAAGGACCGCCTATCTATTACGAATGGAAGGTG
CCCTCGAACGGTCTTGAGCCTCCCAAATTAGACCCACCCATTGGTGTGGATGGACG
TGAGTATCCCGAGACCACTGGAGACTACGGGGTCACCAGCAAGCAGGATGTATTCA
ACACCCGACTAAACGATATTGGAAGCCATCAAAAGAAGCCAGTCCAGATAACCTCG
CCGCTCCAACAGTCGAGCACAAGCCATCGTTTGGCCATCTCGAGATCCATTAAGCC
CAAAGAGGAGCAGGAATCGGCACAGCGGCGATCCGATGTGGTGGCCAGCTCCAC
GGATATAAGCCATCTGCGCAAGCAATTCCTCATTCCGGAGTACGCCTTCCCGCTGG
AAACCATTGGGCGCACGGGTTATGGTCCTGGTGCAGGAGCAGCGGCTGGGGGCT
CCGGCTCCAGCAACGGCGATCTGTATAACTCGTTCCAGCTGAAGATCCCCGAGCA
GCGCGCTAAGTGGTTCGGAGAGAACCCCAAGTGCCCGGAGTGCCATCCTTCGTTC
GTTCTGCCGGGCACCTGCGAGCCCTGCCTGCGCAGATAGggatctttgtgaaggaaccttactt
ctgtggtgtgacataattggacaaactacctacagagatttaaagctctaaggtaaatataaaatttttaagtgtataatgtgtt
aaactactgattctaattgtttgtgtattttagattccaacctatggaactgatgaatgggagcagtggtggaatgccttaatga
gaaaccgtgtccgaag

*UAS-js-dsRED*

In the sequences below, small letters are pUAST sequence. LARGE LETTERS ARE *JS* CDNA SEQUENCE. Underlined letters are dsRED sequence.

agctaaacaatctgcagtaaagtgcaagttaaagtgaatcaattaaaagtaaccagcaaccaagtaaatcaactgcaac
tactgaaatctgccaagaagtaattattgaatacaagaagagaactctgaatagggaattgggaattcgttaacagatctg
smaATGTTGGCACAAACGCAAATTATCCCAACGCCTTGGCGACTTTTGACCCTGGCT
TTATTCTGCTCAGCTTTATTTTTGGCCAGTGAGGCTCAAAATGCCATCCCAAAAACG
AATCCTGCCTCGCTGCTGATACAGAAGACATCCTTCTCCTGCGCCGGACGTCCAGC
CGGATATTATGCGGATGTGGAGACGGGCTGCCAGGTGTACCACATGTGCGATGGC
CTGGGTCGCCAGTTCAGCTACACCTGCCCAAACACGACACTTTTCCAGCAGCGAAT
GCTTATCTGCGACCACTGGTACATGGTGAACTGCTCCAAGGCGGAGAGCAACTATG
CTGCCAATCTCCTAATTGGTCAGCGGGACAAGCCCTTCGTAAACGACGAGGAAAAC
AGCTTGCGCACTCCAAGACCCGATCTTCTGGATCGTCCTTATGCGCCCGACTATTC
CGGCGAGTCCTTCAGAAGCCAATATAAGCAGTTTACTTCCAACCAGAATCAGATACG
TGATGAGTCCGTCAAAGGAGCCGGTGCGGGTAAATCGGATCCCCAGATATCGCAG
ACGCGTTGGCGCATTCCACCACCCAGCCGGACGATCCTTCCACCGGCCTATGAAC
CGCAAATCGAGCTGCCCAGTGCCCAATCGGCCAAGCCCAGGATACCCATTATCACT
AGCACCACCACAACCACTCGAGCGACCACCACCACCCGACCAACAACCACAACTC
GTGCCACCACCACAACAACAACCACTCGAAGACCTCCGGTAACCGCCAGGCCGAA
GGAGGCCTTGCACAACAGGCGGCCAAATTTCCAGGAGCATGATATGGATGACTTGG
GCACCAGCCACAGCACACGGTACAACACCTCGGCAGACTTCAACTCGGCGGAGTC
GCCGCTCCGTGAAACGAAACAGTCCAGCACCAAGCTGACCAAGTTCATCAAGCCG
CCGTCCAAGATCTATGAGCCACCCTTTGTGTACCCCATCTACAATCTAGAAGAGTCC
CAGACCCAGAACGCGGCAGTCGCCACCACACTGCGCACATCAACGGCGGCTCCT
TTCAGTCCTGTTCCAAGTCGCAAGGAGGTCAGCACCACAACTCCACGACCACTGA
GTCGACCGACGACCTTGGCCGGTGTACCCTTCTCCTCTGCCACACACCCAACTAC
GGTTACGACGGTCGGAGTACCGCCACGCAGCGATAATCGGACTCCAGCTCCGGCC
GAGAGCTTCCGCCTGGCCACGCCCACCAGCACTGCAGCACCACCATCGCGCCCG
GCACAGTTGCCCTTCAACGATTTACTGCCGCCGTTCGTTGACTTTGTGCCCCACGA
TATAGCCACCACCCAAGGACCGCCTATCTATTACGAATGGAAGGTGCCCTCGAACG
GTCTTGAGCCTCCCAAATTAGACCCACCCATTGGTGTGGATGGACGTGAGTATCCC
GAGACCACTGGAGACTACGGGGTCACCAGCAAGCAGGATGTATTCAACACCCGAC
TAAACGATATTGGAAGCCATCAAAAGAAGCCAGTCCAGATAACCTCGCCGCTCCAA
CAGTCGAGCACAAGCCATCGTTTGGCCATCTCGAGATCCATTAAGCCCAAAGAGGA
GCAGGAATCGGCACAGCGGCGATCCGATGTGGTGGCCAGCTCCACGGATATAAGC
CATCTGCGCAAGCAATTCCTCATTCCGGAGTACGCCTTCCCGCTGGAAACCATTGG
GCGCACGGGTTATGGTCCTGGTGCAGGAGCAGCGGCTGGGGGCTCCGGCTCCAG
CAACGGCGATCTGTATAACTCGTTCCAGCTGAAGATCCCCGAGCAGCGCGCTAAGT
GGTTCGGAGAGAACCCCAAGTGCCCGGAGTGCCATCCTTCGTTCGTTCTGCCGGG
CACCTGCGAGCCCTGCCTGCGCAGAGGCGGAGGGAGGTCTTCCAAGAATGTTATC
AAGGAGTTCATGAGGTTTAAGGTTCGCATGGAAGGAACGGTCAATGGGCACGAGTT

TGAAATAGAAGGCGAAGGAGAGGGGAGGCCATACGAAGGCCACAATACCGTAAAG
CTTAAGGTAACCAAGGGGGGACCTTTGCCATTTGCTTGGGATATTTTGTCACCACAA
TTTCAGTATGGAAGCAAGGTATATGTCAAGCACCCTGCCGACATACCAGACTATAAA
AAGCTGTCATTTCCTGAAGGATTTAAATGGGAAAGGGTCATGAACTTTGAAGACGGT
GGCGTCGTTACTGTAACCCAGGATTCCAGTTTGCAGGATGGCTGTTTCATCTACAA
GGTCAAGTTCATTGGCGTGAACTTTCCTTCCGATGGACCTGTTATGCAAAAGAAGA
CAATGGGCTGGGAAGCCAGCACTGAGCGTTTGTATCCTCGTGATGGCGTGTTGAA
AGGAGAGATTCATAAGGCTCTGAAGCTGAAAGACGGTGGTCATTACCTAGTTGAATT
CAAAAGTATTTACATGGCAAAGAAGCCTGTGCAGCTACCAGGGTACTACTATGTTGA
CTCCAAACTGGATATAACAAGCCACAACGAAGACTATACAATCGTTGAGCAGTATGA
AAGAACCGAGGGACGCCACCATCTGTTCCTTTAGggatctttgtgaaggaaccttacttctgtggtgt
gacataattggacaaactacctacagagatttaaagctctaaggtaaatataaaatttttaagtgtataatgtgttaaactact
gattctaattgtttgtgtattttagattccaacctatggaactgatgaatgggagcagtggtggaat

*UAS-jsΔCBD-I-dsRED*

In the sequences below, small letters are pUAST sequence. LARGE LETTERS ARE *JS* CDNA SEQUENCE. Underlined letters are dsRED sequence.

agctaaacaatctgcagtaaagtgcaagttaaagtgaatcaattaaaagtaaccagcaaccaagtaaatcaactgcaac
tactgaaatctgccaagaagtaattattgaatacaagaagagaactctgaatagggaattgggaattcgttaacagatctg
smaATGTTGGCACAAACGCAAATTATCCCAACGCCTTGGCGACTTTTGACCCTGGCT
TTATTCTGCTCAGCTTTATTTTTGGCCAGTGAGGCTCAAAATGCCATCCCAAAAACG
AATCCTGCCTCGCTGCTGATACAGAAGACATCCTTCTCCTGCGCCGGACGTCCAGC
CGGATATTATGCGGATGTGGAGACGGGCGGGCGCGCCGATGGCCTGGGTCGCCA
GTTCAGCTACACCTGCCCAAACACGACACTTTTCCAGCAGCGAATGCTTATCTGCG
ACCACTGGTACATGGTGAACTGCTCCAAGGCGGAGAGCAACTATGCTGCCAATCTC
CTAATTGGTCAGCGGGACAAGCCCTTCGTAAACGACGAGGAAAACAGCTTGCGCA
CTCCAAGACCCGATCTTCTGGATCGTCCTTATGCGCCCGACTATTCCGGCGAGTCC
TTCAGAAGCCAATATAAGCAGTTTACTTCCAACCAGAATCAGATACGTGATGAGTCC
GTCAAAGGAGCCGGTGCGGGTAAATCGGATCCCCAGATATCGCAGACGCGTTGGC
GCATTCCACCACCCAGCCGGACGATCCTTCCACCGGCCTATGAACCGCAAATCGA
GCTGCCCAGTGCCCAATCGGCCAAGCCCAGGATACCCATTATCACTAGCACCACCA
CAACCACTCGAGCGACCACCACCACCCGACCAACAACCACAACTCGTGCCACCAC
CACAACAACAACCACTCGAAGACCTCCGGTAACCGCCAGGCCGAAGGAGGCCTTG
CACAACAGGCGGCCAAATTTCCAGGAGCATGATATGGATGACTTGGGCACCAGCCA
CAGCACACGGTACAACACCTCGGCAGACTTCAACTCGGCGGAGTCGCCGCTCCGT
GAAACGAAACAGTCCAGCACCAAGCTGACCAAGTTCATCAAGCCGCCGTCCAAGA
TCTATGAGCCACCCTTTGTGTACCCCATCTACAATCTAGAAGAGTCCCAGACCCAGA
ACGCGGCAGTCGCCACCACACTGCGCACATCAACGGCGGCTCCTTTCAGTCCTGT
TCCAAGTCGCAAGGAGGTCAGCACCACAACTCCACGACCACTGAGTCGACCGACG
ACCTTGGCCGGTGTACCCTTCTCCTCTGCCACACACCCAACTACGGTTACGACGGT
CGGAGTACCGCCACGCAGCGATAATCGGACTCCAGCTCCGGCCGAGAGCTTCCGC
CTGGCCACGCCCACCAGCACTGCAGCACCACCATCGCGCCCGGCACAGTTGCCC

TTCAACGATTTACTGCCGCCGTTCGTTGACTTTGTGCCCCACGATATAGCCACCACC
CAAGGACCGCCTATCTATTACGAATGGAAGGTGCCCTCGAACGGTCTTGAGCCTCC
CAAATTAGACCCACCCATTGGTGTGGATGGACGTGAGTATCCCGAGACCACTGGAG
ACTACGGGGTCACCAGCAAGCAGGATGTATTCAACACCCGACTAAACGATATTGGA
AGCCATCAAAAGAAGCCAGTCCAGATAACCTCGCCGCTCCAACAGTCGAGCACAA
GCCATCGTTTGGCCATCTCGAGATCCATTAAGCCCAAAGAGGAGCAGGAATCGGCA
CAGCGGCGATCCGATGTGGTGGCCAGCTCCACGGATATAAGCCATCTGCGCAAGC
AATTCCTCATTCCGGAGTACGCCTTCCCGCTGGAAACCATTGGGCGCACGGGTTAT
GGTCCTGGTGCAGGAGCAGCGGCTGGGGGCTCCGGCTCCAGCAACGGCGATCTG
TATAACTCGTTCCAGCTGAAGATCCCCGAGCAGCGCGCTAAGTGGTTCGGAGAGAA
CCCCAAGTGCCCGGAGTGCCATCCTTCGTTCGTTCTGCCGGGCACCTGCGAGCCC
TGCCTGCGCAGAGGCGGAGGGAGGTCTTCCAAGAATGTTATCAAGGAGTTCATGA
GGTTTAAGGTTCGCATGGAAGGAACGGTCAATGGGCACGAGTTTGAAATAGAAGGC
GAAGGAGAGGGGAGGCCATACGAAGGCCACAATACCGTAAAGCTTAAGGTAACCA
AGGGGGGACCTTTGCCATTTGCTTGGGATATTTTGTCACCACAATTTCAGTATGGAA
GCAAGGTATATGTCAAGCACCCTGCCGACATACCAGACTATAAAAAGCTGTCATTTC
CTGAAGGATTTAAATGGGAAAGGGTCATGAACTTTGAAGACGGTGGCGTCGTTACT
GTAACCCAGGATTCCAGTTTGCAGGATGGCTGTTTCATCTACAAGGTCAAGTTCATT
GGCGTGAACTTTCCTTCCGATGGACCTGTTATGCAAAAGAAGACAATGGGCTGGGA
AGCCAGCACTGAGCGTTTGTATCCTCGTGATGGCGTGTTGAAAGGAGAGATTCATA
AGGCTCTGAAGCTGAAAGACGGTGGTCATTACCTAGTTGAATTCAAAAGTATTTACA
TGGCAAAGAAGCCTGTGCAGCTACCAGGGTACTACTATGTTGACTCCAAACTGGAT
ATAACAAGCCACAACGAAGACTATACAATCGTTGAGCAGTATGAAAGAACCGAGGG
ACGCCACCATCTGTTCCTTTAGggatctttgtgaaggaaccttacttctgtggtgtgacataattggacaaact
acctacagagatttaaagctctaaggtaaatataaaatttttaagtgtataatgtgttaaactactgattctaattgtttgtgtatttt
agattccaacctatggaactgatgaatgggagcagtggtggaat

*UAS-js∆CBD-II-dsRED*

In the sequences below, small letters are pUAST sequence. LARGE LETTERS ARE *JS* CDNA SEQUENCE. Underlined letters are dsRED sequence.

agctaaacaatctgcagtaaagtgcaagttaaagtgaatcaattaaaagtaaccagcaaccaagtaaatcaactgcaac
tactgaaatctgccaagaagtaattattgaatacaagaagagaactctgaatagggaattgggaattcgttaacagatctg
smaATGTTGGCACAAACGCAAATTATCCCAACGCCTTGGCGACTTTTGACCCTGGCT
TTATTCTGCTCAGCTTTATTTTTGGCCAGTGAGGCTCAAAATGCCATCCCAAAAACG
AATCCTGCCTCGCTGCTGATACAGAAGACATCCTTCTCCTGCGCCGGACGTCCAGC
CGGATATTATGCGGATGTGGAGACGGGCTGCCAGGTGTACCACATGTGCGATGGC
CTGGGTCGCCAGTTCAGCTACACCTGCCCAAACACGACACTTTTCCAGCAGCGAAT
GCTTATCGGGCGCGCCTCCAAGGCGGAGAGCAACTATGCTGCCAATCTCCTAATTG
GTCAGCGGGACAAGCCCTTCGTAAACGACGAGGAAAACAGCTTGCGCACTCCAAG
ACCCGATCTTCTGGATCGTCCTTATGCGCCCGACTATTCCGGCGAGTCCTTCAGAA
GCCAATATAAGCAGTTTACTTCCAACCAGAATCAGATACGTGATGAGTCCGTCAAAG
GAGCCGGTGCGGGTAAATCGGATCCCCAGATATCGCAGACGCGTTGGCGCATTCC

ACCACCCAGCCGGACGATCCTTCCACCGGCCTATGAACCGCAAATCGAGCTGCCC
AGTGCCCAATCGGCCAAGCCCAGGATACCCATTATCACTAGCACCACCACAACCAC
TCGAGCGACCACCACCACCCGACCAACAACCACAACTCGTGCCACCACCACAACA
ACAACCACTCGAAGACCTCCGGTAACCGCCAGGCCGAAGGAGGCCTTGCACAACA
GGCGGCCAAATTTCCAGGAGCATGATATGGATGACTTGGGCACCAGCCACAGCAC
ACGGTACAACACCTCGGCAGACTTCAACTCGGCGGAGTCGCCGCTCCGTGAAACG
AAACAGTCCAGCACCAAGCTGACCAAGTTCATCAAGCCGCCGTCCAAGATCTATGA
GCCACCCTTTGTGTACCCCATCTACAATCTAGAAGAGTCCCAGACCCAGAACGCGG
CAGTCGCCACCACACTGCGCACATCAACGGCGGCTCCTTTCAGTCCTGTTCCAAG
TCGCAAGGAGGTCAGCACCACAACTCCACGACCACTGAGTCGACCGACGACCTTG
GCCGGTGTACCCTTCTCCTCTGCCACACACCCAACTACGGTTACGACGGTCGGAG
TACCGCCACGCAGCGATAATCGGACTCCAGCTCCGGCCGAGAGCTTCCGCCTGGC
CACGCCCACCAGCACTGCAGCACCACCATCGCGCCCGGCACAGTTGCCCTTCAAC
GATTTACTGCCGCCGTTCGTTGACTTTGTGCCCCACGATATAGCCACCACCCAAGG
ACCGCCTATCTATTACGAATGGAAGGTGCCCTCGAACGGTCTTGAGCCTCCCAAAT
TAGACCCACCCATTGGTGTGGATGGACGTGAGTATCCCGAGACCACTGGAGACTAC
GGGGTCACCAGCAAGCAGGATGTATTCAACACCCGACTAAACGATATTGGAAGCCA
TCAAAAGAAGCCAGTCCAGATAACCTCGCCGCTCCAACAGTCGAGCACAAGCCATC
GTTTGGCCATCTCGAGATCCATTAAGCCCAAAGAGGAGCAGGAATCGGCACAGCG
GCGATCCGATGTGGTGGCCAGCTCCACGGATATAAGCCATCTGCGCAAGCAATTCC
TCATTCCGGAGTACGCCTTCCCGCTGGAAACCATTGGGCGCACGGGTTATGGTCCT
GGTGCAGGAGCAGCGGCTGGGGGCTCCGGCTCCAGCAACGGCGATCTGTATAAC
TCGTTCCAGCTGAAGATCCCCGAGCAGCGCGCTAAGTGGTTCGGAGAGAACCCCA
AGTGCCCGGAGTGCCATCCTTCGTTCGTTCTGCCGGGCACCTGCGAGCCCTGCCT
GCGCAGAGGCGGAGGGAGGTCTTCCAAGAATGTTATCAAGGAGTTCATGAGGTTTA
AGGTTCGCATGGAAGGAACGGTCAATGGGCACGAGTTTGAAATAGAAGGCGAAGG
AGAGGGGAGGCCATACGAAGGCCACAATACCGTAAAGCTTAAGGTAACCAAGGGG
GGACCTTTGCCATTTGCTTGGGATATTTTGTCACCACAATTTCAGTATGGAAGCAAG
GTATATGTCAAGCACCCTGCCGACATACCAGACTATAAAAAGCTGTCATTTCCTGAA
GGATTTAAATGGGAAAGGGTCATGAACTTTGAAGACGGTGGCGTCGTTACTGTAAC
CCAGGATTCCAGTTTGCAGGATGGCTGTTTCATCTACAAGGTCAAGTTCATTGGCGT
GAACTTTCCTTCCGATGGACCTGTTATGCAAAAGAAGACAATGGGCTGGGAAGCCA
GCACTGAGCGTTTGTATCCTCGTGATGGCGTGTTGAAAGGAGAGATTCATAAGGCT
CTGAAGCTGAAAGACGGTGGTCATTACCTAGTTGAATTCAAAAGTATTTACATGGCA
AAGAAGCCTGTGCAGCTACCAGGGTACTACTATGTTGACTCCAAACTGGATATAACA
AGCCACAACGAAGACTATACAATCGTTGAGCAGTATGAAAGAACCGAGGGACGCCA
CCATCTGTTCCTTTAGggatctttgtgaaggaaccttacttctgtggtgtgacataattggacaaactacctacaga
gatttaaagctctaaggtaaatataaaatttttaagtgtataatgtgttaaactactgattctaattgtttgtgtattttagattccaa
cctatggaactgatgaatgggagcagtggtggaat
